# Multi-omics reveals mechanisms behind pathogen inhibition by a microalgal microbiome

**DOI:** 10.64898/2026.05.18.725389

**Authors:** Dóra Smahajcsik, Robert A. Koetsier, Emmanuel Tope Oluwabusola, Matilde Emídio Almeida, Line Roager, Scott A. Jarmusch, Morten Dencker Schostag, Joseph Nesme, Marcel Jaspars, Lone Gram, Marnix H. Medema

**Author notes:** Corresponding authors: Lone Gram, Marnix H. Medema. contributed equally (Robert A. Koetsier, Emmanuel Tope Oluwabusola).

## Abstract

Aquaculture is an essential food production sector for meeting the global demand for high-quality protein. However, the sector faces significant challenges from bacterial pathogens, particularly *Vibrio anguillarum*, which causes vibriosis in numerous commercially important fish species. Current disease management strategies rely heavily on antibiotics, leading to antimicrobial resistance and environmental concerns. Microalgal microbiomes represent promising alternatives for sustainable pathogen control, yet the molecular mechanisms underlying their inhibitory activity remain poorly understood. Here, we employed an integrated multi-omics approach to elucidate the mechanisms by which the microbiome of microalga *Isochrysis galbana* inhibits the highly virulent fish pathogen *V. anguillarum* strain 90-11-286. Using a GFP-based inhibition assay, we confirmed potent pathogen suppression by the algal microbiome, achieving complete inhibition at a 1:1000 ratio of pathogen to microbiome. Through 16S rRNA gene amplicon sequencing, metagenomics, metatranscriptomics, and metabolomics, we characterized community composition, genomic potential, gene expression patterns, and metabolite production during pathogen challenge. The inhibitory microbiome was dominated by *Alteromonas macleodii* and *Vreelandella alkaliphila*, with high-quality metagenome-assembled genomes revealing substantial secondary metabolite biosynthetic potential. Metatranscriptomic analysis revealed active expression of biosynthetic gene clusters encoding, for example, non-ribosomal peptide synthetases, particularly a siderophore gene cluster in *V. alkaliphila*. Metabolomic profiling confirmed the production of hydroxamate siderophores in the microbiome, including desferrioxamine analogues, proferrioxamine G1t, and tenacibactin D, which accumulated during pathogen inhibition, as well as 10 putative new compounds. Notably, siderophore production was constitutive rather than pathogen-induced, suggesting iron competition as the primary inhibitory mechanism. Our findings demonstrate that iron sequestration through siderophore production represents a key strategy for pathogen suppression in marine microbial communities. This work provides molecular evidence for microbiome-mediated disease control and establishes a foundation for developing rationally designed multi-strain probiotic consortia for sustainable aquaculture applications, offering an environmentally friendly alternative to antibiotic-based pathogen management strategies.

## INTRODUCTION

In the evolutional arms race between microbial species, secondary metabolites serve as a sophisticated chemical arsenal that mediate microbial interactions and shape community dynamics (Kramer *et al*., 2020; Scherlach and Hertweck, 2021; Chevrette *et al*., 2022; Bech *et al*., 2024). An important class of microbial secondary metabolites are siderophores, specialized iron-chelating compounds that function primarily to sequester this essential element under iron-limited conditions (e.g. marine environments) (Guerinot, 1994; Andrews *et al*., 2003; Miethke and Marahiel, 2007). However, their ecological roles extend far beyond simple resource acquisition: siderophores can serve as signaling molecules, modulate gene expression, virulence factors, and mediate interspecies interactions; serving as a prime example of the intricate chemical crosstalk through which microorganisms navigate cooperative and competitive relationships (Kramer *et al*., 2020). For instance, siderophores can act as a “public good” that can be shared with neighboring microorganisms in a cooperative manner, or as potent inhibitors of competitors by depleting iron from the environment (D’Onofrio *et al*., 2010; Kramer *et al*., 2020). The ecological roles of siderophore production become particularly important in microbial communities in marine environments, where iron availability is limited (Johnson *et al*., 1997). This siderophore-mediated iron acquisition strategy represents a sophisticated form of resource management that has proven effective in terrestrial plant protection (Kloepper *et al*., 1980a, 1980b; Yu *et al*., 2011), and may be even more relevant in marine environments, where iron concentrations are orders of magnitude lower (Johnson *et al*., 1997), suggesting particular potential for microbiome engineering in applied contexts such as disease management in aquaculture (Pybus *et al*., 1994; Gatesoupe, 1997).

Aquaculture is the fastest growing food production sector globally, providing approximately 50% of fish for human consumption and it is projected to play an even more critical role as wild fisheries reach their sustainable limits (FAO, 2022). Despite this growth, intensive rearing of fish and shellfish increases the risks of bacterial disease outbreaks (FAO, 2022). Among bacterial pathogens, members of the genus *Vibrio* are particularly problematic in aquaculture, with *Vibrio anguillarum* recognized as a primary causative agent of vibriosis, a hemorrhagic septicemia affecting over 50 commercially important fish species (Frans *et al*., 2011; Mohamad *et al*., 2019). Current disease management strategies in aquaculture rely heavily on antibiotics, vaccination, and chemical disinfectants (Ringø *et al*., 2014; Cain, 2022). However, these approaches face increasing challenges, such as the development of antimicrobial resistance (AMR), environmental persistence of chemicals, and regulatory restrictions limiting antibiotic use. Vaccination, while highly efficient for adult fish, remains largely unavailable for immunologically immature fish larvae and shellfish (Ringø *et al*., 2014). Thus, there is an urgent need for supplementary sustainable disease control strategies in aquaculture.

Microbial communities are essential to the health and stability of aquatic ecosystems, yet the intricate mechanisms governing their interactions, particularly in the context of pathogen control, remain largely underexplored. Microalgae, as fundamental components of aquatic food webs and common live feed organisms in aquaculture, harbor specialized microbial communities that have co-evolved within the specific ecological niche of the phycosphere (Duerr *et al*., 1998; Natrah *et al*., 2014; Seymour *et al*., 2017). These indigenous microbial communities represent an ecologically relevant reservoir of potential biocontrol agents, likely offering more stable and effective pathogen protection through complex synergistic interactions compared to conventional single-strain probiotics derived from unrelated environments. Their ecological relevance is further enhanced by natural adaptation to the aquaculture food chain, increasing the likelihood of successful establishment within production systems.

While previous investigations have demonstrated that microbiomes of *Isochrysis galbana* can synergistically inhibit the growth of *V. anguillarum* (Smahajcsik *et al*., 2025; Almeida *et al*., 2026), the molecular mechanisms underlying this inhibition remain poorly characterized. Single-strain approaches fail to capture the emergent properties of more complex microbiomes, where community dynamics may facilitate pathogen inhibition through mechanisms not observed in isolation (Timmerman *et al*., 2004; Bertrand *et al*., 2014; Selegato and Castro-Gamboa, 2023).

The “omics-era”, marked by the rapid advancement and decreasing cost of high-throughput sequencing platforms and advanced analytical chemistry has enabled unprecedented insights into complex biological systems through integrated multi-omics approaches (Medema *et al*., 2011; Land *et al*., 2015; Machado *et al*., 2015; Jensen, 2016; Baltz, 2017; Van Der Hooft *et al*., 2020; Blin *et al*., 2023). By simultaneously characterizing community composition, gene expression patterns, and metabolic profiles, we can develop a holistic understanding of the functional attributes driving microbiome-mediated pathogen inhibition. This integrated approach is particularly valuable for investigating microbial communities where functional redundancy, metabolic handoffs, and context-dependent gene expression collectively contribute to emergent properties not predictable from genomic information alone.

In this study, using our previously established GFP-based screening assay (Smahajcsik *et al*., 2025), we identified microalgal microbiomes with inhibitory activity against highly virulent *V. anguillarum* (strain 90-11-286) and subsequently employed a multi-omics approach to elucidate the mechanisms underlying this pathogen inhibition. Through integration of 16S rRNA gene amplicon sequencing, metagenomics, metatranscriptomics, and metabolomics of the inhibitory algal microbiome, we characterized community composition, genetic potential, differential gene expression, and secondary metabolite production associated with inhibitory activity over a time-course of pathogen challenge. Our findings reveal an inhibitory microbiome largely dominated by *Alteromonas macleodii* and *Vreelandella alkaliphila.* We demonstrate the active expression of specific BGCs, including those for siderophore (desferrioxamine) biosynthesis, and link this to the production of a cluster of hydroxamate siderophores, suggesting iron competition as a primary mechanism of pathogen inhibition in this system. Notably, the expression of these key inhibitory pathways appears to be an innate feature of the microbiome rather than a direct, induced response to the pathogen. This work provides a comprehensive understanding of a pathogen inhibitory algal microbiome, offering critical insights into the chemical ecology of pathogen suppression and a foundation for developing rationally designed multi-strain probiotic consortia for sustainable aquaculture.

## MATERIALS AND METHODS

### Bacterial and algal strains and culture conditions

The highly virulent fish pathogen strain, *Vibrio anguillarum* 90-11-286 (Skov *et al*., 1995; Rønneseth *et al*., 2017), carrying a chromosomally integrated GFP construct (pNQFlaC4-gfp27) (Croxatto *et al*., 2007; Almeida *et al*., 2026), hereafter referred to as *V. anguillarum* 90-11-286_gfp, was used as a target pathogen in the inhibition assays. Bacterial strains were stored at −80°C and cultivated in IOCG medium (3% (w/v) Instant Ocean, Aquarium Systems; 0.3% (w/v) HEPES buffer, Thermo Fisher Scientific, CAT No. 11470395; 0.3% (w/v) casamino acids, Thermo Fisher Scientific, CAT No. 223050; 0.2% (w/v) glucose, Merck, CAT No. G8270-100G) at 25°C with shaking at 200 rpm, or on Marine Agar (MA; Difco 2216, BD) at 25°C.

Xenic culture of *Isochrysis galbana* was kindly provided by an aquaculture partner (December 2023). Algal cultures were maintained in f/2 medium with no silicate (f/2 – Si, NCMA); (Guillard and Ryther, 1962; Guillard, 1975) prepared with 3% Instant Ocean (IO; Aquarium Systems Inc., Sarrebourg, France), at 18°C under constant illumination (∼50 μE m^−2^ s^−1^). Cultures were regularly maintained by subculturing (transferring 1% v/v outgrown culture to fresh media) every 4 weeks. Precultures for experiments were initiated as 10% (v/v) subcultures and grown for 5 days prior to experiments.

### Experimental design for GFP-based pathogen inhibition assay and sampling

To investigate the mechanisms of pathogen inhibition by the *I. galbana* microbiome, a GFP-based inhibition assay was conducted in a 96-well plate format as established previously (Smahajcsik *et al*., 2025). *I. galbana* was grown as 10% (v/v) subcultures (from 25 mL to 200 mL in f/2-Si) in four biological replicates for five days. Cultures were then harvested and processed into two fractions: ‘Full Culture’ (FC), containing both algal cells and their associated microbiome, and ‘Filtered Microbiome’ (FM), where algal cells were removed by filtration through a 3 µm pore-size filter. Both fractions (derived from 100 mL of culture each) were concentrated 100-fold by centrifugation (2,445 X *g*, 10 min, 20°C) and resuspension in 1 mL of f/2 −Si medium. The up-concentrated algal samples were stored at 10°C until the next day, when the assay plates were inoculated. On the day of the experiment, serial 10-fold dilutions of these concentrated fractions were prepared. Samples were preserved from each algal culture (1 mL), representing the native microbiome, as well as each up-concentrated fraction (FC-bf and FM-bf; 500 µL) and stored at −80°C until further sample processing.

Overnight culture of *V. anguillarum* 90-11-286_gfp grown in IOCG medium was measured for optical density at 600 nm (OD_600_) and subsequently 10-fold serially diluted to target starting concentrations of 2 log CFU/mL and 4 log CFU/mL. Assay plates (sterile black 96-well plates with flat, clear bottom; Perkin Elmer 6005430) were inoculated with combinations of algal fraction dilutions (FC and FM, 10^−1^, 10^−2^, 10^−3^ and 10^−4^) and the pathogen at either concentration (180 µL total well content), in four biological replicates under the following experimental conditions: microbiomes with pathogen present (VFC, VFM), microbiomes without pathogen (FC, FM), *Vibrio* monocultures (V) and sterile media control (C). The pathogen inhibition assay was conducted for 26 hours at 25°C with continuous monitoring every 20 min using a Perkin Elmer EnVision 2104 Multilabel reader. GFP fluorescence (excitation 485 nm, emission 535 nm, gain = 100) was measured as an indicator of pathogen growth, alongside optical density at 620 nm. Each measurement cycle included 20 s orbital shaking prior to detection. Sample preparation and the inhibition assay are visualized in Figure 1 (left panel).

**Figure 1.**
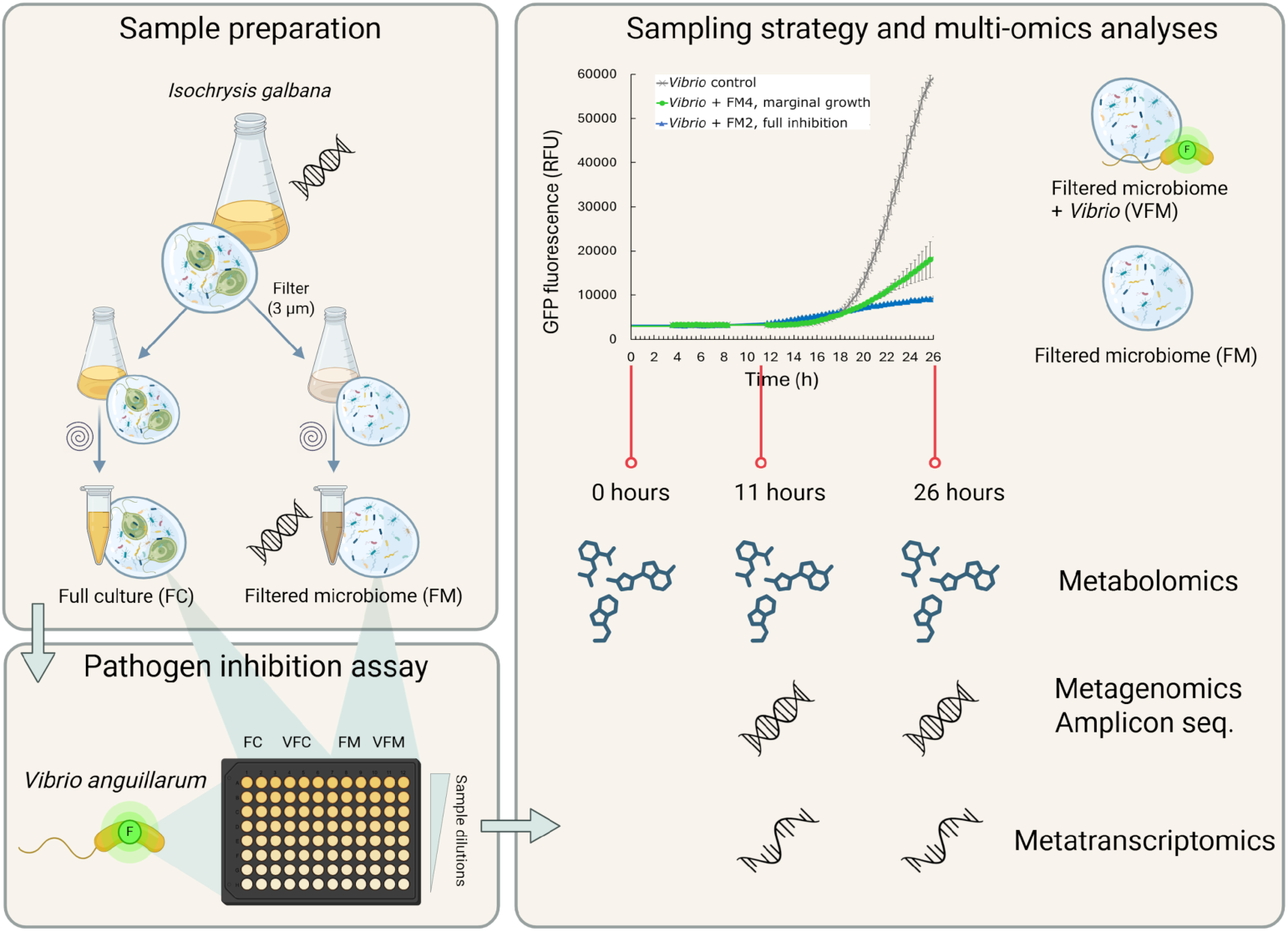
Experimental design and sampling strategy for the multi-omics analyses. The microbiome of microalga *Isochrysis galbana* was processed into two up-concentrated fractions (top left panel): the full culture (FC), containing both algal and bacterial cells, and the filtered microbiome (FM) with only bacterial cells. The pathogen inhibition of FC and FM fractions was tested in the GFP-based inhibition assay against GFP-tagged highly virulent *Vibrio anguillarum* strain 90-11-286 (bottom left panel). Samples were taken at 0 hours, 11 hours and 26 hours after co-inoculation (right panel). Samples showing full inhibition (FM2) or allowing marginal growth of the pathogen (FM4) of the microbiome cultured with or without pathogen (VFM and FM, respectively) were processed for multi-omics analyses: metabolomics (0-hour, 11-hour, 26-hour timepoints); metagenomics (11-hour, 26-hour timepoints); metataxonomic through amplicon sequencing (11-hour, 26-hour timepoints and pre-assay samples FM and the native microbiome); and metatranscriptomics (11-hour, 26-hour timepoints). Figure created with BioRender.com.

Samples for multi-omics analyses (metagenomics, metatranscriptomics, metabolomics, and 16S rRNA gene amplicon sequencing) were collected at three time points: t = 0, t = 11, and t = 26 hours post-inoculation, representing different phases of interaction and inhibition. Each experimental condition consisted of four biological replicates, with three technical replicates per sample which were pooled prior to extraction. To facilitate sampling and minimize handling time, each biological replicate was set up in a separate plate and was inoculated in three technical replicates (allowing to harvest the entire plate at each sampling time), resulting in a total of 12 assay plates.

At each sampling timepoint, a batch of plates was removed from the incubator and 20 µL ice-cold RNA stop solution (5% phenol in absolute ethanol) (Feike *et al*., 2012) was added immediately to each well and mixed. The 96-well plates were centrifuged (2,250 X *g*, 15 min, 20°C), and the supernatant was transferred to sterile, round bottom plastic 96-well plates. The pellets (for nucleic acid extraction) as well as the supernatants (for metabolite extraction) were stored at −80°C until further sample processing. The sample strategy for the multi-omics analyses is summarized in Figure 1 (right panel).

### DNA and RNA co-extraction

Since inhibition profiles of FC and FM samples were comparable, only FM samples were processed to focus on the microalgal microbiome. Total DNA and RNA were co-extracted from FM samples collected at the 11- and 26-hour timepoints, from wells exhibiting complete inhibition of the pathogen (VFM2) and marginal pathogen growth (VFM4), alongside corresponding microbiome dilutions without pathogen inoculation (FM2 and FM4, respectively). Samples of medium sterility control (C) and *Vibrio* monocultures (V) were also included. Native algal microbiomes and up-concentrated FM-bf samples collected prior to the inhibition assays were also processed. All samples included four biological replicates, resulting in a total of 56 samples.

DNA and RNA co-extraction was performed using the RNeasy® PowerMicrobiome® Kit (Qiagen, CAT No. 26000-50) according to the manufacturer’s protocol. All surfaces and equipment were UV-sterilized and treated with RNase*Zap*™ (Invitrogen, CAT No. AM9780) prior to extraction. All consumables were certified RNase/ DNase free. Keeping samples on ice, the pellets from three technical replicates of each sample were resuspended in PM1-ß-mercaptoethanol buffer (included in the kit), pooled and transferred to 1.5 mL bead beat tubes (included in the kit). Following elution in 100 µL of elution buffer (included in the kit), the eluate was split into two 50 µL fractions for separate DNA and RNA downstream processing. Samples were stored at −80°C in smaller aliquots.

For RNA samples (FM, VFM, V, C samples collected at 11 and 26 hours), contaminating DNA was removed using the TURBO DNA-*free*™ Kit (Invitrogen, CAT No. AM1907) following the manufacturer’s instructions. RNA quality and integrity was assessed using the RNA 6000 Nano Kit and a 2100 Agilent Bioanalyzer (Agilent Technologies), and concentration was determined using a Qubit Fluorometer with the HS RNA assay kit (Invitrogen).

For DNA samples, RNA was digested using RNase A (ThermoFisher, DNase and protease-free, 10 mg/mL, CAT No. EN0531). DNA quality and purity was assessed using a DeNovix 439 DS-11+ spectrophotometer (DeNovix Inc., Wilmington, DE, USA) and confirmed by agarose gel electrophoresis. DNA concentrations were measured using a Qubit 2.0 Fluorometer with the Qubit dsDNA HS Assay Kit (Invitrogen).

### 16S rRNA gene amplicon sequencing and analysis

The V3-V4 hypervariable regions of the 16S rRNA gene were amplified from a total of 50 DNA samples (FM, VFM, V, C samples collected at 11 and 26 hours, as well as prior-assay samples: raw and FM-bf) using primers 341f (5’-CCTACGGGNGGCWGCAG-3’) and 805r (5’-GACTACHVGGGTATCTAATCC-3’) carrying unique octametric barcodes for sample multiplexing (Klindworth *et al*., 2013) (Table S1). PCR products were purified using Agencourt AMPure XP magnetic beads (0.8:1 bead volume to DNA solution; Agencourt Bioscience Corporation, Beverly, MA, USA). Concentrations of the PCR products were measured using a Qubit 2.0 Fluorometer with the high sensitivity (HS) DNA assay kit (Qubit dsDNA HS assay; Invitrogen by Thermo Fisher Scientific Inc., Eugene, OR, USA) and pooled in equimolar concentrations (target 250 ng DNA per sample). Sequencing was performed on an Illumina NovaSeq 6000 platform generating paired-end 250-bp reads by Novogene (Cambridge, United Kingdom).

Bioinformatic processing of 16S rRNA gene amplicon data was performed using QIIME 2 (v2024.10.1) (Bolyen *et al*., 2019). After importing the raw sequences, the pipeline began with demultiplexing and primer/barcode removal via the cutadapt demux-paired plugin (Martin, 2011). Quality filtering, sequence denoising, and dereplicating were performed using DADA2 (Callahan *et al*., 2016), and an amplicon sequence variant (ASV) table was generated. Phylogenetic relationships were inferred using MAFFT v.7.525 (Katoh and Standley, 2013) and a phylogenetic tree was constructed with FastTree v2.2 (Price *et al*., 2010). Taxonomic assignment was conducted using a custom classifier trained on the Silva reference database (version 138.2) with the study-specific primer sequences, implemented through the feature-classifier plugin (Werner *et al*., 2012; Bokulich *et al*., 2018). ASVs lacking phylum-level classification or identified as chloroplast/mitochondrial sequences were subsequently removed from the dataset.

### Long-read metagenomic sequencing and analysis

A pooled DNA sample, representative of the inhibitory *I. galbana* microbiome across inhibitory conditions (equimolar pooling of all 26 h samples, including FM2, VFM2, FM4 and VFM4, to a total of 1 µg DNA), was prepared for long-read sequencing. Library preparation was performed using the Ligation Sequencing Kit (SQK-LSK114, Oxford Nanopore Technologies) according to the manufacturer’s protocol, opting for the Long Fragment Buffer to enrich DNA fragments of 3 kb or longer. Sequencing was conducted on a single MinION flow cell (FLO-MIN114, Oxford Nanopore Technologies), loading a total of 300 ng DNA with the library.

The minimum read length filter was set to 1 kb and raw reads were basecalled using Dorado (v7.4.14) with the super-accurate basecalling (v4.3.0), 400 bps setting. Reads shorter than 1,000 bp were removed using Filtlong (v0.2.1) (Wick, 2025a). Adapters and barcodes were trimmed using Porechop (0.2.4) (Wick, 2025b) with default settings, including analysis of 10,000 reads for adapter definition and chimera removal. Quality checks before and after trimming were performed using NanoPlot (v1.42.0) (De Coster and Rademakers, 2023) and FastQC (v0.12.1) (Andrews, 2010). Metagenome assembly was performed using metaFlye (v2.9.5, Python v3.12.8) (Kolmogorov *et al*., 2020). The assembly was first polished using Medaka (v2.0.1) (nanoporetech/medaka, 2025) with default settings. A second polishing step was performed with Pilon (v1.24) (Walker *et al*., 2014), using the short-read metatranscriptomics dataset, with the –fix snps option to correct only single base errors. Binning of contigs into Metagenome-Assembled

Genomes (MAGs) was performed using Metabat2 (v2.17) (Kang *et al*., 2019). SAM and BAM file manipulations were carried out using Samtools (v1.21) (Danecek *et al*., 2021), and bin quality was assessed using CheckM2 (v1.1.0). Taxonomic assignment of the final MAGs was performed using GTDB-Tk (v2.4.0) (Chaumeil *et al*., 2022; Parks *et al*., 2022) and bin6 and bin7 were taxonomically assigned with CAT_pack (-bins option) (v6.0.1) (von Meijenfeldt *et al*., 2019). Functional annotation of MAGs was performed using Bakta (v1.11) (Schwengers *et al*., 2021). Secondary metabolite biosynthetic gene cluster (BGC) prediction and analysis was conducted using antiSMASH (v7.1.0) (Blin *et al*., 2023) in loose mode with additional features including MIBiG database comparison, ClusterBlast, and KnownClusterBlast analyses. Assembly statistics were assessed using Quast (v5.3.0) (Mikheenko *et al*., 2023) before and after polishing steps.

### Metatranscriptomic sequencing and analysis

RNA samples were pre-processed as described above. Quality control removed samples with apparent contamination (one replicate of each sample from 11 hour time-point). Furthermore, all samples with marginal pathogen growth (FM4, VFM4) were excluded due to insufficient *Vibrio* RNA yield (even in monoculture) for meaningful analysis, resulting in 22 samples being shipped to Novogene (Cambridge, United Kingdom). Four additional samples were excluded due to insufficient RNA concentrations, leaving 18 samples for processing. Novogene performed rRNA depletion, RNA fragmentation, and reverse transcription to cDNA prior to library preparation. Strand-specific sequencing was performed on Illumina NovaSeq X Plus Series (PE150) generating paired-end 150-bp reads.

Raw metatranscriptomic reads were processed using fastp (v0.24.0) (Chen, 2025) for quality control and adapter trimming, with quality assessed by FastQC (v0.12.1) (Andrews, 2010) before and after. Trimmed reads from all samples were mapped to the polished metagenomic assembly (generated as described above) using Bowtie2 (v2.5.4) (Langmead *et al*., 2019). The resulting SAM files were converted to BAM format, sorted, and indexed using Samtools (v1.21) (Danecek *et al*., 2021). Gene expression levels were quantified using featureCounts (v2.0.8) (Liao *et al*., 2014).

Differential gene expression analysis was performed using the R package DESeq2 (v1.48.1) (Love *et al*., 2014), with lfcThreshold = 0, altHypothesis = “greaterAbs”, and a significance level of alpha = 0.05. An additive model was utilized to assess statistical significance, comparing expression levels at 26 h versus 11 h while averaging expression across the FM and VFM communities, and vice versa. P-values were generated using the Wald test and subsequently adjusted for multiple testing using the Benjamini-Hochberg false discovery rate correction (Benjamini and Hochberg, 1995).

The gene expression counts from RNA-seq read mapping and quantification were subjected to further processing for co-expression analysis in R v4.1.1 (R Core Team, 2023). Only genes that were found in one of the previously created taxonomic bins were considered for co-expression analysis. To reduce noise, genes that did not pass a threshold of at least 10 counts in half of the samples were excluded from further analysis. For this filtering step, only samples in which a gene-carrying organism was estimated to be present (>35% of its genes detected) were considered. Organisms for which more than half of the genes were removed in low expression filtering were not included in co-expression analysis, to prevent misinterpretation of their transcriptional activity. The expression of the remaining genes was normalized for each taxon individually (Klingenberg and Meinicke, 2017), using the trimmed mean of M-values from ‘edgeR’ v3.42.2 (Robinson and Oshlack, 2010). Subsequently, the normalized expression values were transformed using hyperbolic arcsine transformation (Johnson and Krishnan, 2022). To prevent an excess of high correlation values,confounding factors were corrected for by removing the first principal component from the data in a further processing step (Parsana *et al*., 2019).

Co-expression analysis was conducted for the remaining taxa individually to study associations in BGC activity and to learn about specialized metabolic activity of the community members. Information on BGC regions was extracted from the ‘.json’ summary output file of antiSMASH (v7.1.0). BGCs were subdivided into co-expressed subclusters. For subcluster creation, hierarchical clustering was conducted using 1-Pearson’s correlation as a distance metric with ‘ward.D2’ linkage. The hierarchical clustering tree was cut at h=0.5 using the ‘stats::cutree’ function to create BGC subclusters. Similarity in gene expression between BGC subclusters was quantified using Spearman’s correlation. For this, all genes of BGC subclusters were correlated, and the mean of these values was calculated after Fisher z-transformation to prevent introduction of bias in this step (Alexander, 1990).

To assess potential relationships between incomplete bins and high-quality MAGs, bin-level correlation analysis was performed using Pearson’s correlation coefficient to determine the association between the total counts (sum of all gene expression counts) of the bins, correlating across all samples. This analysis helped identify whether incomplete bins represented genomic fragments or plasmids associated with the larger MAGs.

### Chemical extraction of samples for untargeted metabolomics

All samples (assay supernatants) from each sampling timepoint (0 h, 11 h, 26 h; a total of 72 samples) were processed for metabolomic analysis. Technical triplicates of each sample were pooled to 1 mL deep well plates and dried using a cooled vacuum centrifuge. Subsequently, 600 µL acetonitrile was added to each sample, and the plate was ultrasonicated for 15 minutes and dried under nitrogen flow. To remove contaminants and salts, samples were resuspended in methanol and processed using C18 Spin Tips (100 µL bed, Thermo Scientific Pierce, Product code: 10615555). Spin tips were preconditioned with methanol, washed with water, and metabolites were eluted with methanol. Eluted samples were dried under nitrogen flow and stored at −20°C until reconstitution and LC-MS analysis. Subsequent untargeted metabolomic analysis included four biological replicates of the following sample types: sterile medium control (C), *Vibrio* monoculture (V), microbiome with complete pathogen inhibition (VFM2) and marginal pathogen growth (VFM4), and corresponding microbiome dilutions without pathogen inoculation (FM2 and FM4).

### Untargeted metabolomics analysis

Extracts were analyzed using a ThermoScientific Orbitrap IQ-X MS coupled to a Thermo Instruments UPLC system under the following operating conditions: capillary temperature 275°C, auxiliary gas flow rate 10 arbitrary units, sheath gas flow rate 50 arbitrary units, static gas mode, sweet gas (Arb) 1.0, ion transfer tube temperature 275°C, vaporizer temperature 350°C, and capillary positive ion spray voltage 3.5k, and a lock mass correction set at RunStart EASY-IC. The full MS was set with a scan mass range of 150-2000 m/z, Orbitrap resolution at 120000, RF Lens at 35%, absolute AGC target at 4.000E5, maximum injection time mode at Auto, and data type set to centroid with the data-dependent acquisition mode set at Cycle Time (Top Speed). The assisted HCD (high-energy collisional dissociation) energy at 30% with an activation time of 10 milliseconds and the Orbitrap resolution at 60000 were set for the MS/MS fragmentation step.

A stationary column of the LC system consists of a diode array detector (DAD) with a variable wavelength detection range of 200-800 nm, an analytical Kinetex 2.6 µm EVO C18 column (100 Å, 100 × 2.1 mm), and a mobile phase gradient of 0.1% formic acid in LC-MS grade H₂O: acetonitrile from 5–95% in 13 minutes, and isocratic elution at 95% for 5 minutes at a flow rate of 300 µL/min with MS acquisition for 17 minutes was employed in the UPLC and MS system.

### Data processing and molecular annotation of metabolomics data

A molecular network was created with the Feature-Based Molecular Networking (FBMN) (Nothias *et al*., 2020) on Global Natural Product Social Molecular Networking (GNPS) (Wang *et al*., 2016) from the Orbitrap IQ-X mass spectrometry data. The raw MS datasets were centroided through conversion to mzML format using MSConvert from the ProteoWizard (Chambers *et al*., 2012). The mzML files were further pre-processed with MZmine version 4.5.0 (Schmid *et al*., 2023) to generate the final datasets including quantitative MS1 file and MGF files of fragment ions required for the FBMN analysis by adopting the following settings parameter: mass ions detection was achieved through the removal of the baseline noise level of the MS1 and MS2 ions set to 1.0 ×10^4^ and 1.0 ×10^2^, respectively. The chromatograms construction for the MS1 ions were detected with the ADAP chromatogram builder where minimum consecutive scans, minimum intensity for consecutive scans, minimum absolute height, m/z mass tolerance (scan-to-scan) were set to 4, 1.0 × 10^2^ and 1.0 × 10^4^ and 5 ppm, respectively. In order to create the chromatogram of each mass ion while maintaining the initial feature list generated in the preceding stage, the peak detection employed the local minimum resolver as a deconvolution technique. The chromatographic threshold for this algorithm was set at 85%, the minimum search range in the RT absolute range was set at 0.05 min, the minimum absolute height was set at 2.0E6, the minimum relative height was set at 0.0%, and the minimum ratio of peak top to edge was set at 1.70. The identified peaks were deisotoped using isotopic peaks grouper in which the m/z tolerance (intra-sample) was set to 5 ppm, the RT tolerance to 0.02 for absolute minutes, maximum charge of 3 absolute minutes, and the representative isotope as the most intense. The deisotoped peaks were subsequently aligned using a join aligner to correct any deviation of the retention time of each mass peak: the ion m/z tolerance for the alignment was set to 10 ppm, retention time tolerance to 0.06 at absolute minute, weight for m/z and RT set to 75 and 25, respectively, and the mobility weight was set to 1. The peak list produced after the alignment step was filtered using a feature list row filter algorithm to remove any mass ions outside the set range of 100 to 2000 m/z and keep peak rows that match all criteria.

The resulting feature table results were exported to the GNPS platform for FBMN analysis. The data was filtered by removing all MS/MS fragment ions within +/− 17 Da of the precursor m/z. MS/MS spectra were window filtered by choosing only the top 6 fragment ions in the +/− 50 Da window throughout the spectrum. The precursor ion and MS/MS fragment ion mass tolerances were 0.05 Da and 0.05 Da. A molecular network was then created where edges were filtered to have a cosine score above 0.70 and more than 6 matched peaks. Further, edges between two nodes were kept in the network if and only if each of the nodes appeared in each other’s respective top 20 most similar nodes. Finally, the maximum size of a molecular family was set to 100, and the lowest-scoring edges were removed from molecular families until the molecular family size was below this threshold. The spectra in the network were then searched against GNPS spectral libraries. The library spectra were filtered in the same manner as the input data. All matches kept between network spectra and library spectra were required to have a score above 0.7 and at least 6 matched peaks. The molecular networks were visualized using Cytoscape 3.10 software (Shannon *et al*., 2003).

The mzML files from MSConvert were imported to the SIRIUS software (v4.2.0) (Dührkop *et al*., 2019) for chemical dereplication, a bioinformatic framework readily available for putative annotation of compound structures and molecular formula prediction of mass spectrometry data using CSI-FingerID percentage scoring to predict molecular fingerprints afforded by the MS/MS features of the high-resolution acquisition MS data of complex compound mixtures (Dührkop *et al*., 2015). In addition to the CSI:Finger ID, the introduction of COSMIC (Compound Structure Identification by Mass Spectrometry and In Silico Chemistry) brought about a high level of increased confidence in the structural annotation pipeline (Hoffmann *et al*., 2022). Upon importing the files into SIRIUS, an automatic profiling of the precursor ions and molecular adducts displayed on the platform was filtered, ordering according to their retention time or molecular mass. The m/z belonging to the blank control were manually removed, and interesting mass ions were computed using the following general setting parameters: the instrument was set to Orbitrap, filtered by isotope pattern, MS2 mass accuracy set to 10 ppm, and candidates of predicted molecular formula set to 10 with a minimum of 1 candidate per ion stored, while other general settings were set as default. The C, H, O, N, S, and P were selected at auto-detection. The CSI: Fingerprint Prediction, including Predict FPS for putative adducts, score threshold, and database search (Bio Database, GNPS, PubChem, COCONUT, and Natural Product), was checked. The results of each dataset, including the molecular formula, mass error ppm, and CSI-FingerID matching scores of the predicted structures recorded.

## RESULTS

### Inhibition of *Vibrio anguillarum* 90-11-286 by the *Isochrysis galbana* microbiome

To assess the pathogen-inhibitory potential of the *I. galbana* microbiome, we employed a GFP-based assay to monitor the growth of *V. anguillarum* 90-11-286_gfp. The initial *V. anguillarum* inoculum was standardized from an overnight culture with an optical density corresponding to 8 log CFU/mL, diluted to target concentrations of 2 and 4 log CFU/mL for the assay. Plate counts confirmed 1.9 and 3.9 CFU/mL in the starting inocula.

The algal cultures had an algal density of 5.10 (± 0.06) log cells/mL and the filtered microbiome (FM) fraction, used for detailed downstream analysis, had a starting bacterial concentration of 7.5 (± 0.26) log CFU/mL after up-concentration, while the bacterial density in the up-concentrated full culture (FC) fraction was 7.9 (± 0.20) log CFU/mL.

The *I. galbana* microbiome effectively inhibited *V. anguillarum* growth (Figure 2). Complete inhibition, defined as conditions where GFP fluorescence showed no exponential increase (<17% of the maximum control signal) after 26 hours, was achieved at a 1:1000 ratio of *V. anguillarum* to the algal microbiome (FC or FM) at the lower *Vibrio* inoculation density (2 log CFU/mL) (Figure S1A). At the higher *Vibrio* inoculation density (4 log CFU/mL), the same ratio reduced GFP fluorescence to 24% (FC) and 27% (FM) of the maximum control signal after 26 hours (Figure S1B). The inhibition profiles between FC and FM fractions were comparable, thus subsequent analyses focused on the FM samples to specifically investigate the role of the microalgal bacterial community, minimizing interference from the alga itself. To elucidate the mechanisms underlying pathogen inhibition, FM samples at the lower *Vibrio* inoculation density (2 log CFU/mL) were selected for downstream analyses: FM2, representing complete inhibition at a 1:1000 ratio (10⁻² dilution), and FM4, representing marginal *Vibrio* growth at a 1:10 ratio (10⁻⁴ dilution) (Figure 2).

**Figure 2.**
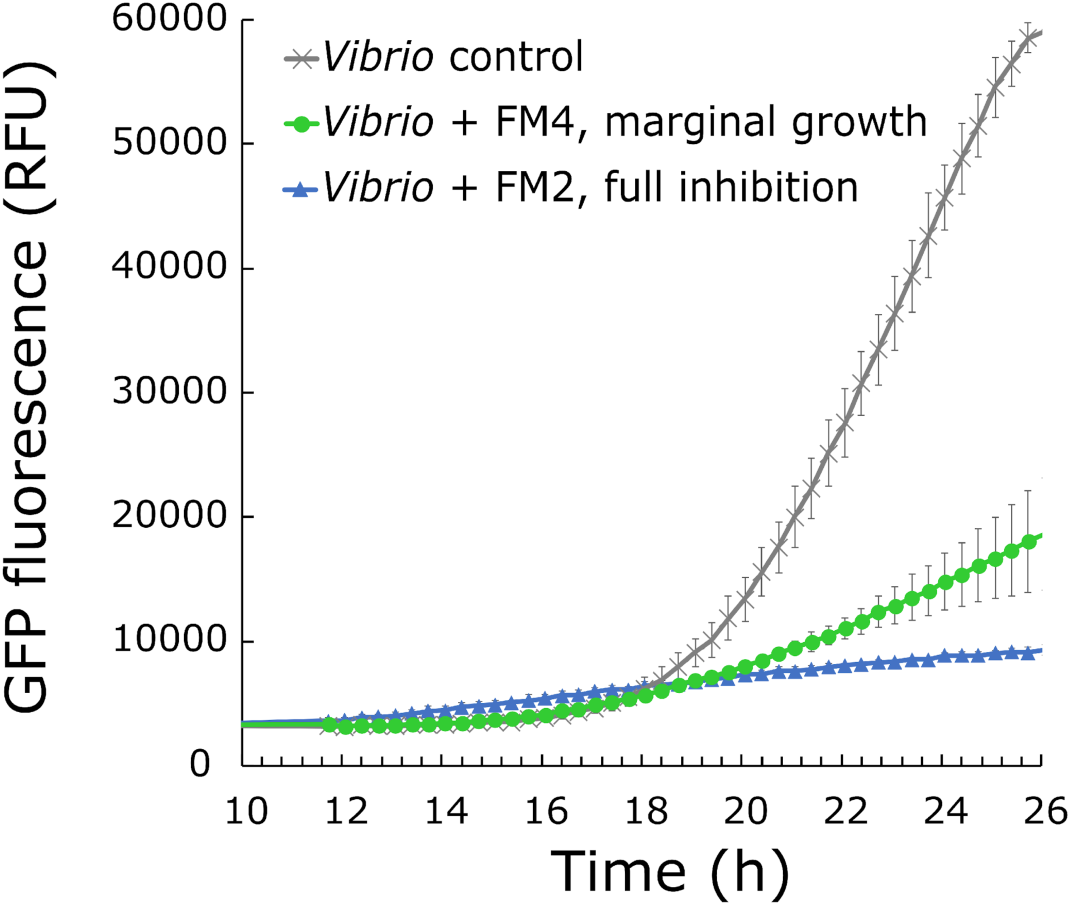
Inhibition of *V. anguillarum* 90-11-286_gfp by the filtered microbiome (FM) of *Isochrysis galbana* during a 26-hour co-cultivation, as shown by decreased GFP signal compared to *Vibrio* control (2 log CFU/mL starting concentration; n=3). Full inhibition of the pathogen was achieved by 1:1000 ratio of *V. anguillarum*: FM (10^−2^ dilution of FM; *Vibrio* + FM2; n=4), whereas a 1:10 ratio allowed marginal growth of the pathogen (10^−4^ dilution of FM; *Vibrio* + FM4; n=4). Error bars represent standard deviation.

### Microbiome composition and dynamics during pathogen challenge

To characterize the composition of the *I. galbana* microbiome and its response to pathogen challenge, 16S rRNA gene amplicon sequencing was performed on 50 samples collected at 11, and 26 hours from samples with full inhibition (FM2 and VFM2) and marginal pathogen growth (FM4 and VFM4). The sequencing yielded a total of 73,590,344 raw reads, with over 93% of reads having a Q30 score and over 98% a Q20 score, averaging 1,471,806 reads per sample. After bioinformatic processing, including denoising, chimera removal, and filtering of chloroplast/mitochondrial sequences and amplicon sequencing variants (ASVs) not assigned to a phylum, 327 ASVs remained, comprising 8,332,506 reads, with a mean read count of 166,650 per sample.

The native *I. galbana* microbiome was dominated by genera including *Muricauda*, *Alteromonas*, *Winogradskyella*, *Sulfitobacter*, *Halomonas*, *Roseovarius*, *Nisaea*, *Lentilitoribacter*, *Parasphingorhabdus*, and an unclassified member of the *Paracoccaceae* family (formerly *Rhodobacteraceae*) (Figure 3A). Upon up-concentration and filtration (FM-bf samples), the taxonomic profile showed shifts in relative abundances, notably a reduction in *Winogradskyella* and the unclassified *Paracoccaceae* member.

**Figure 3.**
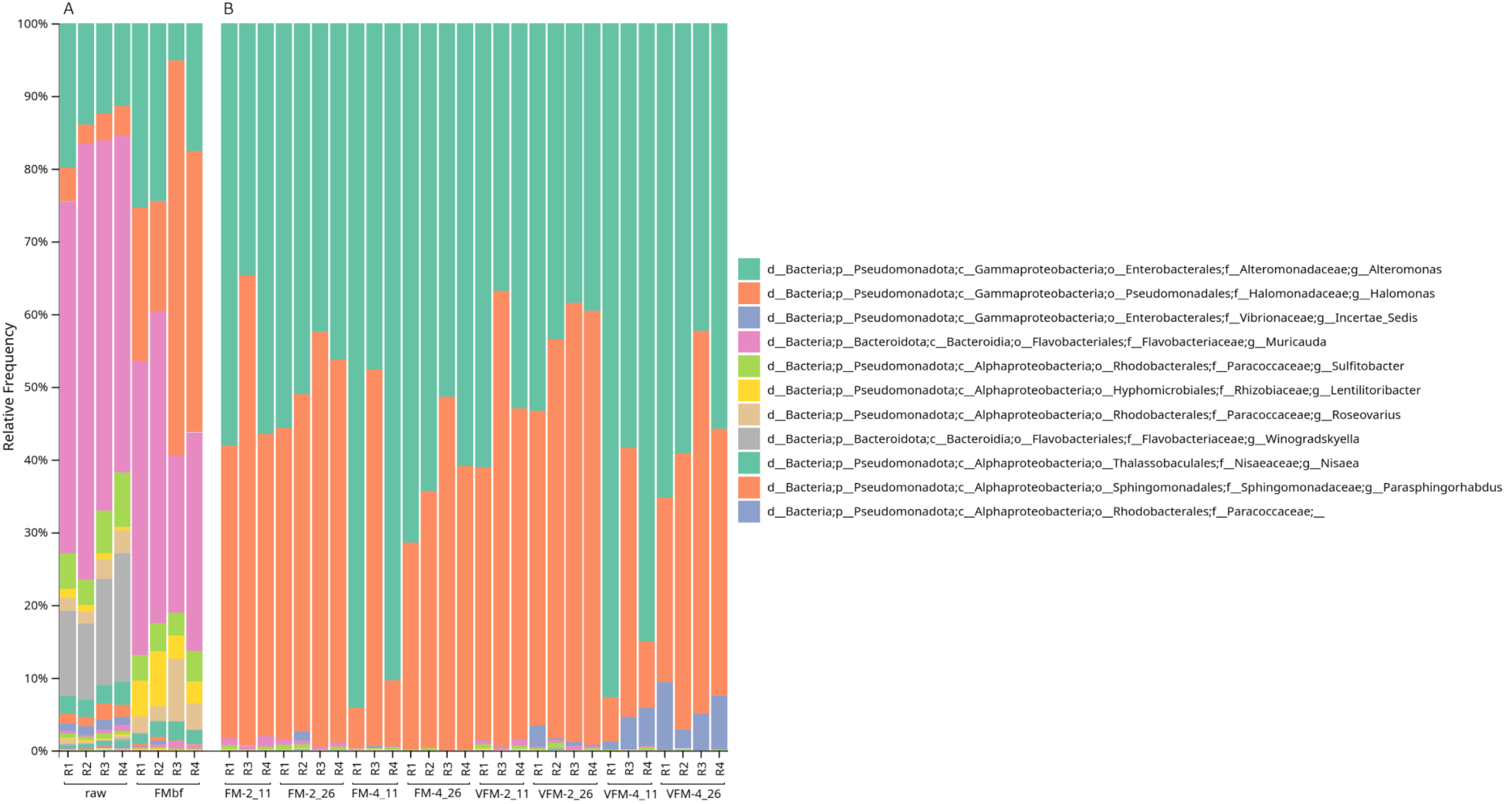
Microbial composition of the *Isochrysis galbana* microbiome before (**A**), and after the *Vibrio*-inhibition assay (**B**). Native microbiome (raw) and up-concentrated filtered-microbiome (FMbf) represent the algal microbiome prior to the assay (**A**). The *Vibrio*-inhibitory microbiome was analyzed after 11-hours and 26-hours of incubation, with and without pathogen inoculation (VFM and FM, respectively), representing a fully inhibitory microbiome (FM2) and a partially inhibitory microbiome, allowing marginal growth of the pathogen (FM4) (**B**). The inhibitory microbiome is less diverse and is largely dominated by *Alteromonas* and *Halomonas* (by new classification: *Vreelandella*) genera, with higher relative abundance of *Vibrio* in the VFM4 samples (marginal pathogen growth).

During the pathogen inhibition assay, the community structure of the inhibitory microbiomes (e.g., VFM2 conditions) was primarily composed of *Alteromonas*, *Halomonas* (later identified as *Vreelandella*), *Muricauda*, and *Sulfitobacter* (Figure 3B). In samples exhibiting only marginal pathogen growth (VFM4), the target pathogen *Vibrio* was also detected at higher relative abundances. Notably, *Vreelandella alkaliphila* (previously classified as *Halomonas alkaliphila*) and *Alteromonas macleodii* emerged as key constituents of the inhibitory microbiome, alongside *Sulfitobacter* spp.

### Metagenomic assembly reveals genomic potential of key community members

To investigate the genomic potential of the inhibitory microbiome, pooled DNA from inhibitory FM samples at t=11 h was subjected to long-read Nanopore sequencing. This generated 4.88 million reads with an estimated N50 of 10.32 kb and 33.13 Gb of estimated bases. After basecalling (100% efficiency, 3.76 million reads >Q10, 30.6 Gb), adapter removal and further quality filtering, 3.01 million reads (27.4 Gb, 80.1% of initial Q10 data) were used for assembly. These processed reads had a mean length of 9,098 bp, an N50 of 10,721 bp, and a mean quality score of 18.7, with 93.4% of reads exceeding Q15.

Metagenome assembly using MetaFlye yielded 132 contigs, totaling 22.677 Mb, with an N50 of 3.127 Mb and a largest contig of 4.73 Mb. The assembly showed high contiguity, with 112 contigs ≥10 kb. Polishing with Medaka and Pilon (using short-read metatranscriptomic data, 98.96% overall alignment rate) resulted in a final assembly of 22.674 Mb with no ambiguous bases and a GC content of 49.29%.

Metagenomic binning with MetaBat2 recovered eight bins. Five of these were classified as high-quality metagenome-assembled genomes (MAGs) and three low-completeness bins (Table 1). High ANI values (96.35-99.63%) to reference genomes confirmed species-level assignments for the high-quality MAGs and bin1, and two low-completeness bins (bin.6 *Alteromonas*, bin.7 Rhodobacterales), though at higher taxonomic levels (Table 1).

**Table 1.** Overview of metagenome-assembled genomes (MAGs) and low-completeness bins recovered from the inhibitory I. galbana microbiomes. Metagenomic assembly resulted in three complete genomes (MAG1, MAG2 and MAG3), two high quality drafts (MAG4 and MAG5), and three low-completeness bins (bin1, bin6 and bin7). Taxonomic classification of the five high quality MAGs and bin1 was based on ANI, using GTDB-Tk, while bin6 and bin7 were classified with CAT_pack and showed lower confidence levels.

| Bin/<br>MAG | Taxonomy<br>(Family/ Genus/<br>Species) | Confidence<br>(%) | Completeness<br>(%) | Contamination<br>(%) | Size (Mb) | Average<br>coverage |
| --- | --- | --- | --- | --- | --- | --- |
| bin3<br>(MAG1) | *Alteromonadaceae*/<br>*Alteromonas*/<br>*Alteromonas macleodii* | 97.32 | 100 | 0.06 | 4.73<br>single contig | 2781x |
| bin5<br>(MAG2) | *Halomonadaceae*/<br>*Vreelandella*/<br>*Vreelandella alkaliphila* | 98.6 | 100 | 0.22 | 4.36<br>single contig | 1650x |
| bin8<br>(MAG3) | *Vibrionaceae*/<br>*Vibrio*/<br>*Vibrio anguillarum* | 98.41 | 100 | 0.19 | 4.32<br>in 2 contigs | 56x |
| bin2<br>(MAG4) | *Rhodobacteraceae*/<br>*Sulfitobacter*/<br>*Sulfitobacter pontiacus* | 97.12 | 90.31 | 2.22 | 3.49 | 6.1x |
| bin4<br>(MAG5) | *Flavobacteriaceae*/<br>*Flagellimonas*/<br>*Flagellimonas sp.* | 96.35 | 93.39 | 3.61 | 3.61 | 7x |
| bin1 | *Rhodobacteraceae*/<br>*Phaeobacter*/<br>*Phaeobacter piscinae* | 99.63 | 6.95 | 0.00 | 0.29 | 2.3x |
| bin6 | *Alteromonadaceae*/<br>*Alteromonas* | 35 | 4.34 | 0.03 | 0.26 | 6506x |
| bin7 | Rhodobacterales | 58 | 5.71 | 0.04 | 0.20 | 12.3x |

Functional annotation with Bakta predicted 21,459 coding sequences (CDSs) across the metagenome (coding density 88.8%), including 335 tRNAs, 7 tmRNAs, 73 rRNAs, and 66 ncRNAs. AntiSMASH analysis in loose mode predicted 100 BGCs across these bins (26 in relaxed mode), indicating substantial secondary metabolic potential within the community. Two BGCs from the *Vibrio anguillarum* MAG showed high similarity to MIBiG (Zdouc *et al*., 2025) reference gene clusters for anguibactin and vanchrobactin, two well-characterized siderophores of this species (Actis *et al*., 1986; Soengas *et al*., 2006), with conserved NRPS domain architecture and PARAS-predicted substrate specificities (Terlouw *et al*., 2025) supporting these annotations. Other BGCs involved clusters classified as: NRPS and NRPS-like, betalactone, arylpolyene, RiPP-like, terpene, ranthipeptide, NI-siderophore, T1PKS, T3PKS, hserlactone, and the majority as saccharides and fatty acids.

### Metatranscriptomic analysis reveals active BGC expression in dominant taxa, with particular focus on Vreelandella alkaliphila

To profile gene expression dynamics, metatranscriptomic sequencing was performed on samples from FM2 and VFM2 conditions at 11 h and 26 h. Sequencing yielded between 29.6 and 814.5 million raw reads per sample (mean: 565.7 million), with high quality (mean Q20: 98.39%, mean Q30: 95.89%). After quality filtering and trimming, reads were mapped to the polished metagenomic assembly with an overall mapping rate of 98.96% (93.49% unique alignments). FeatureCounts assigned 80.8% of the total mapped reads (767,480,582 reads) to 21,466 annotated features across 130 contigs, indicating robust coverage of expressed genes within the microbial community. The majority of the transcripts could be mapped back to the two main community members, *Alteromonas macleodii* and *Vreelandella alkaliphila*, with *Vibrio* transcripts detected in meaningful quantities only in *V. anguillarum* monocultures (Figure 4A). Bin-level correlation analysis revealed that two incomplete bins, classified as *Alteromonas* sp. and Rhodobacterales, are associated with the high-quality *Alteromonas macleodii* and *Sulfitobacter pontiacus* MAGs, respectively, suggesting they might represent plasmids, which was confirmed by the presence of characteristic plasmid replication (repA, repB, repC), partitioning (parB), and conjugation (type IV secretion system (virB operon, virD4)) genes in both bins (Figure 4B). While this observation is likely, bins are kept separate for downstream analysis in the absence of confirmation using ie. chromosome-plasmid crosslinking or dedicated assembly methods (Castañeda-Barba *et al*., 2025; Piera Líndez *et al*., 2026).

**Figure 4.**
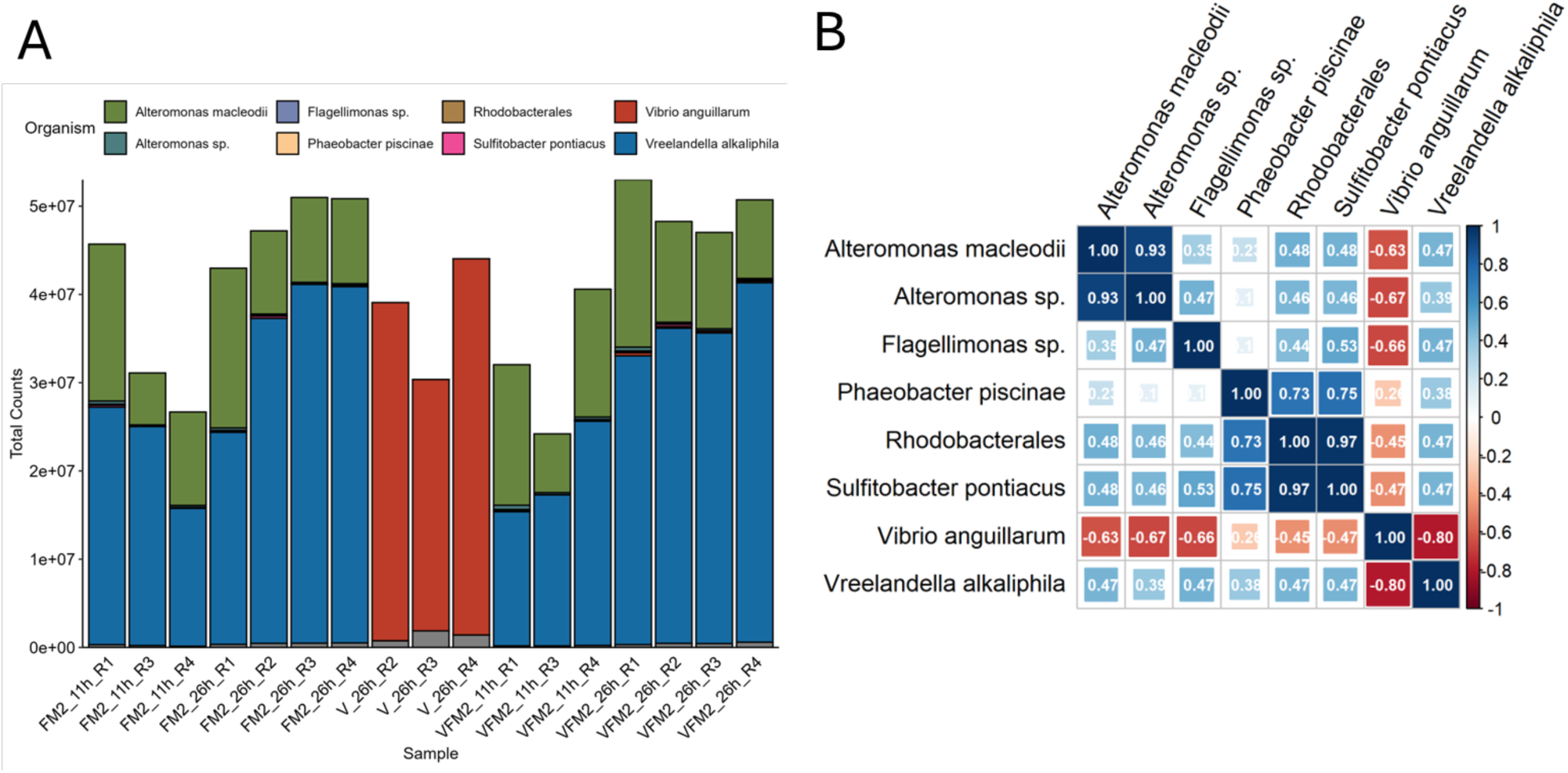
Taxonomic distribution of RNA-seq expression and bin correlations reveal community dynamics and genomic associations. (**A**) Total expression counts per sample. The majority of expression counts originate from the dominant community members *Alteromonas macleodii* and *Vreelandella alkaliphila*. *Vibrio anguillarum* transcripts are only detected in meaningful counts in outgrown pure cultures (after 26 hours). (**B**) Correlation analysis of the bins indicates that incomplete bins classified as *Alteromonas* sp. and Rhodobacterales are related to the *Alteromonas macleodii* and *Sulfitobacter pontiacus* MAGs, respectively, likely representing plasmids.

Investigation of the BGC expression profiles of the dominant community members revealed distinct responses to the experimental conditions. In the *Vreelandella alkaliphila* MAG, expression of BGC_3.7, annotated as a non-ribosomal peptide synthetase (NRPS) cluster (putative NI-siderophore, homologous to desferrioxamine BGC), differed significantly between the two timepoints of the pathogen challenge (26 h vs 11 h; 7 out of 10 genes differentially expressed, median FDR-adjusted p-value: 3.45 × 10⁻¹¹), with higher expression during the 11 h timepoints compared to 26 h, but did not differ based on the presence of the pathogen (FM2 and VFM2) (Figure 5A). Another *V. alkaliphila* BGC, BGC_3.14 (putative betalactone), showed significantly higher expression at 26 h compared to 11 h (16 out of 17 genes differentially expressed, median FDR-adjusted p-value: 1.98 × 10⁻¹³), particularly in the VFM2 – 26 h condition (Figure 5B). The expression of these BGCs appeared to be a constitutive feature of the microbiome rather than an induced response to the pathogen.

**Figure 5.**
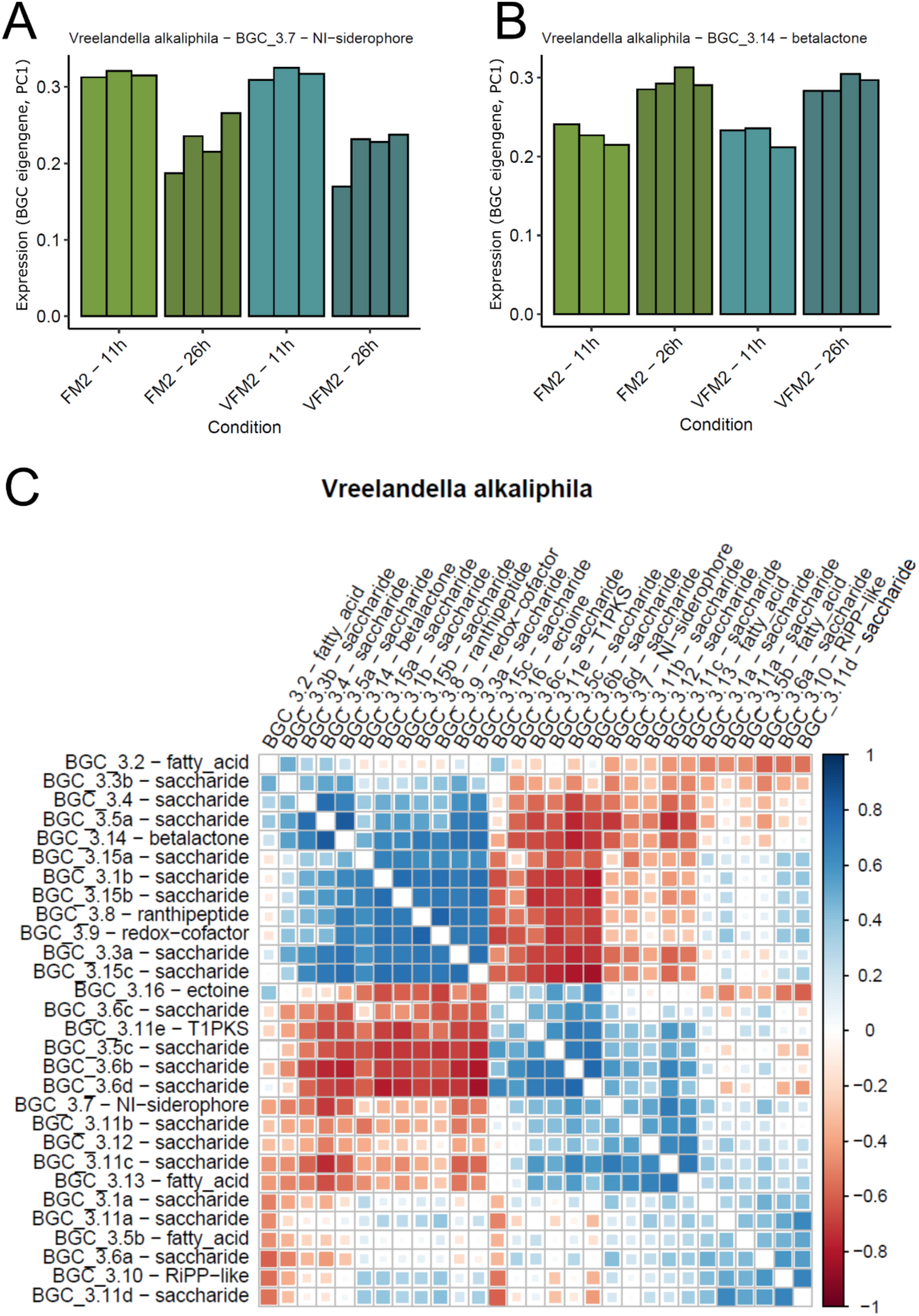
Transcriptional modulation of selected biosynthetic gene clusters (BGCs) with predicted antimicrobial potential (**A**, **B**) and BGC co-expression (**C**) by *Vreelandella alkaliphila*. (**A**) Expression of the NI-siderophore BGC across samples with full pathogen inhibition at different timepoints (11 h and 26 h), representing the inhibitory microbiome with (VFM2) and without (FM2) the pathogen. Expression is higher at 11 h timepoints compared to 26 h timepoints, indicating higher production during the early phase of the pathogen challenge. Expression is unaffected by the presence of the pathogen. (**B**) Expression of the betalactone BGC across samples with full pathogen inhibition at different timepoints (11 h and 26 h), representing the inhibitory microbiome with (VFM2) and without (FM2) the pathogen. Expression is higher at 26 h timepoints compared to 11 h timepoints, suggesting production at the later stage of the pathogen challenge. Expression is unaffected by the presence of the pathogen. (**C**) BGC co-expression by *V. alkaliphila*.

Co-expression analysis of BGCs within *Vreelandella alkaliphila* revealed clusters of BGCs with correlated expression profiles, suggesting coordinated regulation of secondary metabolite production (Figure 5C). For instance, BGC_3.2 (fatty acid), BGC_3.3b (saccharide), BGC_3.4 (saccharide), BGC_3.5a (saccharide) and BGC_3.14 (betalactone) formed a positively correlated cluster, while they showed negative correlation with a group of BGCs including BGC3.11e (T1PKS) and BGC_3.7 (NI-siderophore). Positive correlation of BGC_3.7 (NI-siderophore) was shown with BGC3.11e (T1PKS) and a number of saccharide BGCs (BGCs 3.6c,b,d; 3.5c; 3.11b,c; 3.12) and a fatty acid BGC (BGC_3.13). This positive correlation with a group of saccharide-encoding gene clusters (non-canonical BGCs), including BGC_3.11 which was identified as a putative Wzy-dependent exopolysaccharide biosynthesis cluster by epssmash (Daugberg *et al*., 2025), suggests potential coordinated regulation of biofilm-related polysaccharide production. Ultimately, the co-expression analysis indicates complex regulatory interplay governing the expression of different BGCs, and while these preliminary findings reveal intriguing patterns, it warrants further analysis and deeper interpretation.

### Metabolomic profiling confirms siderophore production during pathogen inhibition

An untargeted metabolomics study using feature-based molecular networking (FBMN) was conducted via LC-MS/MS ESI-Orbitrap analysis on supernatants from six sample types: sterile medium control (C), *Vibrio* monoculture (V), microbiome with complete pathogen inhibition (VFM2), microbiome with marginal pathogen growth (VFM4), and the corresponding microbiome dilutions without pathogen inoculation (FM2 and FM4). Samples were collected at 0 h, 11 h, and 26 h. These conditions were selected for untargeted metabolomic profiling based on bioactivity observed in an initial screen of 72 extracts against *Vibrio anguillarum* 90-11-286. The aim was to characterise the molecular ecosystem and identify metabolites associated with the observed inhibitory activity.

FBMN is a computational approach that uses mass fragmentation data; specifically, m/z values, retention times, and precursor intensities to assess spectral and feature similarity, thereby organising related molecules into clusters. The molecular network analysis revealed 1025 nodes representing molecular entities detected in the VFM4 co-culture metabolome across the three timepoints. After removing features originating from the medium and *Vibrio* monoculture controls, 34 molecular features remained and were organised into clusters and singletons. Detailed inspection of the VFM4 network highlighted a major cluster of molecular ions that emerged prominently at 26 h (Figure 6). Chemical dereplication of this predominant cluster, performed using Thermo Scientific FreeStyle™ 1.8 SP2 QF1 for manual inspection based on accurate mass, molecular formula, and MS/MS fragmentation, and SIRIUS 4.2 for structure prediction (drawing from >110 million structures in databases such as PubChem and ChemSpider) (Brittin *et al*., 2024), led to the putative identification of hydroxamate siderophores produced during the 26-hour incubation (Table 2, Figure 6 and S2).

**Figure 6.**
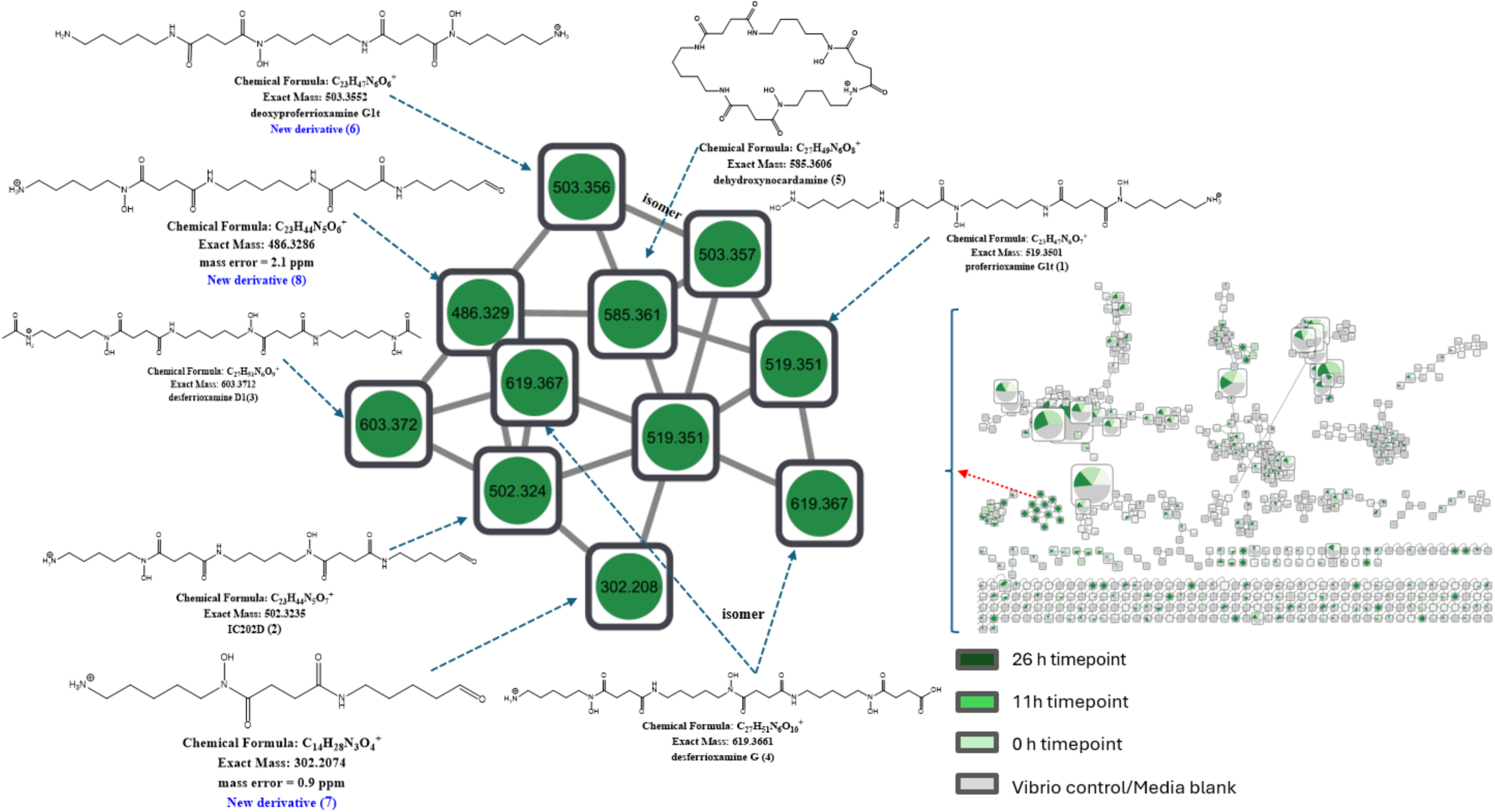
Feature-based Molecular Networking analysis of *Vibrio* + FM4 (VFM4) extracts exhibiting marginal pathogen growth (partial inhibition) at 26 hrs.

**Table 2.** Chemical dereplication of putative hydroxamate siderophores from VFM2 and VFM4 extracts.

| Extracts_type | CSI:Finger ID score | RT [min] | Observed m/z [M+H] | MF[M+H] | Theoretical mass [M+H] | Error ppm | Putative compounds (PubChem, Reaxys.com and NP Atlas) |
| --- | --- | --- | --- | --- | --- | --- | --- |
| V-FM2, V-FM4 |  | 8.2 | 503.3557 | C<sub>23</sub>H<sub>47</sub>N<sub>6</sub>O<sub>6</sub> | 503.3552 | 1.2 | No hit |
| V-FM2, V-FM4 | -32.9 | 8.3 | 519.3504 | C<sub>23</sub>H<sub>47</sub>N<sub>6</sub>O<sub>7</sub> | 519.3501 | 0.4 | Proferrioxamine G1t |
| V-FM2, V-FM4 |  | 8.4 | 302.2077 | C<sub>14</sub>H<sub>28</sub>N<sub>3</sub>O<sub>4</sub> | 302.2074 | 0.9 | No hit |
| V-FM2, V-FM4 |  | 8.7 | 485.3592 | C<sub>26</sub>H<sub>49</sub>N<sub>2</sub>O<sub>6</sub> | 485.3585 | 1.3 | No hit |
| V-FM4 |  | 8.9 | 486.3296 | C<sub>23</sub>H<sub>44</sub>N<sub>5</sub>O<sub>6</sub> | 486.3286 | 1.9 | No hit |
| V-FM2 | -114.0 | 9.1 | 603.4084 | C<sub>28</sub>H<sub>55</sub>N<sub>6</sub>O<sub>8</sub> | 603.4076 | 1.3 | Tenacibactin D |
| V-FM2, V-FM4 | -6.3 | 9.2 | 502.3241 | C<sub>23</sub>H<sub>44</sub>N<sub>5</sub>O<sub>7</sub> | 502.3235 | 1.2 | IC202D |
| V-FM2, V-FM4 | -59.3 | 9.3 | 603.3718 | C<sub>27</sub>H<sub>51</sub>N<sub>6</sub>O<sub>9</sub> | 603.3712 | 1 | Desferrioxamine D1 |
| V-FM2 | -63.8 | 9.3 | 517.3349 | C<sub>23</sub>H<sub>45</sub>N<sub>6</sub>O<sub>7</sub> | 517.3344 | 0.9 | IC202C |
| V-FM2 |  | 9.5 | 625.3145 | C<sub>34</sub>H<sub>41</sub>N<sub>6</sub>O<sub>5</sub> | 625.3133 | 1.9 | No hit |
| V-FM2, V-FM4 | -69.4 | 9.5 | 619.3665 | C<sub>27</sub>H<sub>51</sub>N<sub>6</sub>O<sub>10</sub> | 619.3661 | 0.5 | Desferrioxamine G |
| V-FM2 |  | 9.8 | 585.3975 | C<sub>28</sub>H<sub>53</sub>N<sub>6</sub>O<sub>7</sub> | 585.3970 | 1.3 | No hit |
| V-FM2, V-FM4 |  | 9.9 | 316.2122 | C<sub>16</sub>H<sub>30</sub>NO<sub>5</sub> | 316.2118 | 1.3 | No hit |
| V-FM2 |  | 9.9 | 633.3827 | C<sub>28</sub>H<sub>53</sub>N<sub>6</sub>O<sub>10</sub> | 633.3818 | 1.5 | No hit |
| V-FM2 | -6.3 | 10.3 | 573.3250 | C<sub>25</sub>H<sub>45</sub>N<sub>6</sub>O<sub>9</sub> | 573.3243 | 1.2 | Desferrioxamine X1 |
| V-FM2, V-FM4 | -6.5 | 10.5 | 585.3610 | C<sub>27</sub>H<sub>49</sub>N<sub>6</sub>O<sub>8</sub> | 585.3606 | 0.6 | Dehydroxynocardamine |
| V-FM2 | -2.7 | 10.7 | 601.3565 | C<sub>27</sub>H<sub>49</sub>N<sub>6</sub>O<sub>9</sub> | 601.3555 | 1.6 | Desferrioxamine E |
| V-FM2 |  | 16.8 | 513.2314 | C<sub>21</sub>H<sub>33</sub>N<sub>6</sub>O<sub>9</sub> | 513.2314 | -0.6 | No hit |
| V-FM4 |  | 21.5 | 835.5547 | C<sub>44</sub>H<sub>71</sub>N<sub>10</sub>O<sub>6</sub> | 835.5553 | -0.6 | No hit |

'No hit' represents a putative new molecule that was not found in the Natural product databases. Sirius CSI:Finger ID score = The score is typically negative, and closer to zero means there is a higher likelihood that the molecular ions are the same to the predicted database hit based on MS/MS fragmentation pattern.

Among the putatively identified compounds were molecular ions at *m/z* 519.3504, 502.3241, 603.3718, and 619.3665, corresponding to the linear hydroxamate siderophores proferrioxamine G1t (**1**), IC202D (**2**), desferrioxamine D1 (**3**), and desferrioxamine G (**4**), respectively (Figure S3-S6). A molecular ion at *m/z* 585.3610 (C₂₇H₄₉N₆O₈, Δ = +0.6 ppm), annotated as the cyclic siderophore dehydroxynocardamine (**5**), was also detected. In addition, three further putative new derivatives were dereplicated based on MS/MS fragmentation and FBMN analysis (Figure 6). One newly detected derivative, designated deoxyproferrioxamine G1t (**6**), exhibited a precursor ion at *m/z* 503.3557 (C₂₃H₄₇N₆O₆, Δ = +1.2 ppm) and represents a close analogue of **1**, differing by 16 Da in a manner consistent with terminal deoxygenation (Figure 6 and S8-S9). A second derivative (**7**) at *m/z* 302.2080 (C₁₄H₂₈N₃O₄, Δ = +0.9 ppm) displayed a characteristic 184-Da core fragment shared with **2**, supporting its structural annotation (Figure 6 and S10-S11). A third mass ion at *m/z* 486.3296, which clustered with **2**, showed a diagnostic neutral loss of 16 Da from the hydroxamate N–O adjacent to the carbonyl. This fragmentation pattern, characteristic of hydroxamate cleavage, supports its assignment as a new analogue (**8**) lacking an oxygen atom on the hydroxamate functional group (Figure 6 and S12-13).

Additional clusters containing unique metabolites detected at 11 h in VFM4 were also present at 0 h (Figure 6), indicating that these compounds were not associated with the pathogen inhibition observed during co-culture.

In the VFM2 extracts, which showed complete inhibition of *Vibrio*, the molecular networking analysis revealed 1295 nodes. Most molecular features were shared between the controls and the VFM2 extracts at the 0 h timepoint, with the exception of a single major molecular cluster that appeared at 26 h, mirroring the pattern observed for VFM4 (Figure S18). The putative bioactive metabolites were predominantly annotated as siderophores; however, VFM2 produced a greater diversity and abundance of these compounds, including several derivatives not detected in the VFM4 extracts. These included both linear and cyclic hydroxamate siderophores, such as tenacibactin D (9), IC202C (10), desferrioxamine X1 (11), and desferrioxamine E (12) (Figure S10–S18; Table 2).

To further assess this observation, a semi-quantitative comparison of FBMN precursor node intensities between VFM2 and VFM4 at 26 h was conducted (Figure S19). This analysis showed that VFM2 contained a substantially higher number of siderophore-related nodes than VFM4. Several precursor nodes also exhibited higher intensities in VFM2, indicating greater concentrations of both the shared putative hydroxamate siderophores and the additional unknown metabolites (Figure S3–S18; Table 2). These findings suggest that increased production or accumulation of iron-chelating metabolites under the VFM2 condition contributed to the complete pathogen inhibition, potentially through enhanced iron depletion.

A total of eight putative new metabolites were detected within the siderophore clusters of both VFM2 and VFM4 extracts, along with two additional metabolites from other clusters. Four of these were present exclusively in VFM2 (Table 2). These unique metabolites may also contribute to the observed activity; however, the current data are insufficient to support this hypothesis.

## DISCUSSION

This study employed an integrated multi-omics approach to dissect the mechanisms underlying the inhibition of the fish pathogen *Vibrio anguillarum* by the microbiome associated with the microalga *Isochrysis galbana*. Our findings confirmed that the *I. galbana* microbiome effectively suppresses *V. anguillarum* growth, as previously described in (Smahajcsik *et al*., 2025). The combined insights from 16S rRNA gene sequencing, metagenomics, metatranscriptomics, and metabolomics point towards iron competition, mediated by the production of hydroxamate siderophores, as a primary inhibitory mechanism in the inhibitory microbiome largely dominated by *Vreelandella alkaliphila* and *Alteromonas macleodii*.

The consistent and strong inhibition observed with the FM2 samples (full inhibition of *V. anguillarum*) underscores the potency of the bacterial community in controlling *V. anguillarum*. The dominance of *Alteromonas macleodii* and *Vreelandella alkaliphila* in these inhibitory microbiomes, as revealed by both 16S rRNA profiling and MAG recovery, positions them as key players. *Alteromonas* species are ubiquitous marine bacteria, frequently associated with algal cultures (Natrah *et al*., 2014; Roager *et al*., 2023; Smahajcsik *et al*., 2025). These bacteria demonstrate beneficial interactions with their algal hosts, including growth enhancement of *Isochrysis galbana* (Sandhya and Vijayan, 2019; Cao *et al*., 2021) and production of bioactive compounds such as homoserine lactones, siderophores (Sandhya and Vijayan, 2019; Koch *et al*., 2020) and antibacterial agents against pathogens like *Vibrio harveyi* (Abraham, 2004).

Reflecting this metabolic potential, our metagenomic and metatranscriptomic data also revealed a transcriptionally active homoserine lactone and ribosomally synthesized and post-translationally modified peptide (RiPP) BGC, though the specific functions of these BGCs in our system warrant further investigation.

*Vreelandella alkaliphila* (formerly *Halomonas alkaliphila*) belongs to the *Vreelandella* genus of halotolerant or even halophilic bacteria widely distributed across diverse saline environments, including oceanic niches. Members of the *Halomonas* genus have been described for their metabolic versatility, producing diverse bioactive compounds including hydrolytic enzymes, extracellular polysaccharides (EPS) (Biswas *et al*., 2022), antibiotics (Velmurugan *et al*., 2013), antifungals (Bibi *et al*., 2018) and pigments (Fariq *et al*., 2019; Athmika *et al*., 2021). Consistent with this biosynthetic potential, our metatranscriptomic analysis revealed that *V. alkaliphila* harbors numerous BGCs encoding saccharides, alongside several other potentially bioactive BGCs. Although direct probiotic effects of *V. alkaliphila* remain underexplored, related *Halomonas* species have demonstrated biocontrol activity against aquaculture pathogens, including *Vibrio harveyi* and *Vibrio parahaemolyticus* (Suantika, 2013; Velmurugan *et al*., 2013). Additionally, *Halomonas* species are known to synthesize siderophores such as desferrioxamine (Hintersatz *et al*., 2023), a metabolic potential that aligns with our present findings and complements our ongoing investigation of *Vreelandella alkaliphila* and *Sulfitobacter pontiacus* co-cultures (Smahajcsik *et al*., 2026).

A central finding of this study is the strong evidence for siderophore-mediated iron competition in the algal microbiome, leading to the inhibition of *V. anguillarum*. The metatranscriptomic data revealed active expression of a BGC predicted to synthesize a non-ribosomal peptide (NI-siderophore, homologous to a desferrioxamine BGC) in *Vreelandella alkaliphila*, particularly in the early phase of the pathogen challenge (11 hour timepoint). Notably, the siderophore production genes were co-expressed with saccharide-encoding BGCs, potentially indicating concurrent biofilm formation – a defensive strategy well documented in related *Halomonas* species (Qurashi and Sabri, 2012; Balmaceda *et al*., 2022). The predicted siderophore production was corroborated by the metabolomic analysis, which detected a suite of hydroxamate siderophores, including various desferrioxamine analogues (e.g., D1, E, G, X1), proferrioxamine G1t, tenacibactin D, and dehydroxynocardamine, particularly after 26 hours of incubation (by which time they have accumulated) in the VFM2 samples with complete pathogen inhibition. The higher precursor ion intensities and more derivatives of these siderophores in the fully inhibitory samples (VFM2) compared to the marginally inhibitory samples (VFM4) strongly supports their role in the inhibition. By sequestering iron, they limit its availability to *V. anguillarum*, a pathogen known to heavily rely on iron for growth and virulence (Frans *et al*., 2011). Iron limitation is a well-established strategy for microbial competition in marine environments, where iron is often a limiting nutrient (Gram, 1993; Smith and Davey, 1993; Miethke and Marahiel, 2007; Kramer *et al*., 2020). Hence, our results are in line with previous studies highlighting siderophore production as an important mechanism for pathogen biocontrol in aquaculture as well as plants (Kloepper *et al*., 1980b, 1980a; Gram, 1993; Gatesoupe, 1997; Miethke and Marahiel, 2007), however here for the first time shown as an effect of a marine microbiome consortium.

The active expression of the siderophore BGC in *V. alkaliphila*, even in the absence of the pathogen (FM2 samples), suggests that this iron-scavenging capability is an inherent trait of the microbiome. Rather than representing an induced defense, siderophore production may be essential for microbial persistence within the community, providing a pre-emptive defense mechanism as a secondary effect, by limiting iron availability. This proactive strategy could be ecologically advantageous, allowing the established microbiome to maintain a low-iron environment that is less conducive to the proliferation of invading pathogens like *V. anguillarum*. The co-expression analysis of BGCs within *V. alkaliphila* further indicated a complex regulatory network governing its secondary metabolism. Notably, siderophore production is potentially coordinated with other metabolic functions, such as biosynthetic features encoded by non-canonical BGCs, related to fatty acid and saccharide metabolism. The expression of other BGCs, such as the putative betalactone cluster in *V. alkaliphila*, which showed increased expression at the later phase of the pathogen challenge (26 h timepoint), suggests that additional bioactive compounds might contribute to the overall inhibitory effect or play roles in later stages of microbial interaction, warranting further investigation.

The multi-omics approach was crucial in connecting genomic potential (MAGs and BGCs) to transcriptional activity and, ultimately, to the production of functional metabolites. Each omics layer provided distinct but complementary insights that, together, paint a more nuanced picture of siderophore function in the inhibitory microbiome. Metagenomics revealed the biosynthetic potential encoded in numerous BGCs, while metatranscriptomics showed that siderophore BGC expression was independent of pathogen presence yet significantly higher at the early timepoint (11 h), suggesting that siderophore production is a constitutive trait linked to microbial establishment. Meanwhile, metabolomics revealed higher siderophore accumulation at the later timepoint (26 h), indicating that despite declining transcriptional activity, the functional products continue to accumulate in the environment, increasingly restricting iron availability for invading pathogens. This link is exemplified by the detection of desferrioxamine-like siderophores - well-characterized hydroxamate siderophores (Sandy and Butler, 2009; Jarmusch *et al*., 2021) - in the metabolome, which correspond directly to the NI-siderophore BGC actively expressed in *V. alkaliphila*, confirming the connection between genomic potential, transcriptional activity, and functional metabolite production. Together, these findings suggest that siderophore production is initiated as a constitutive trait during microbial establishment, with the resulting metabolites persisting and accumulating to generate an iron-limited environment that acts as a competitive defense mechanism. This integration provides a more robust understanding than any single omics dataset could offer alone and exemplifies the power of such approaches in microbial ecology.

Our study focused on the filtered microbiome (FM) to specifically elucidate the role of the bacterial community associated with *Isochrysis galbana*. While this approach minimized interference from the alga, it is possible that direct algal contributions or synergistic interactions between the alga and its microbiome in the full culture (FC) could play additional roles in pathogen defense in the natural phycosphere environment. Future studies could explore these interactions in more detail. Furthermore, while several siderophores were putatively identified, we detected 10 compounds in the siderophore cluster with no hits to any Natural product database, indicating putative new compounds. Some other detected metabolites remain uncharacterized and could also contribute to the observed inhibition. *Sulfitobacter pontiacus* represents interesting avenues for future research as a species known to be associated with microalgae (Amin *et al*., 2015). In another study we demonstrated the potential of *Sulfitobacter* to enhance inhibition of *V. anguillarum* when co-cultured with *V. alkaliphila* (Smahajcsik *et al*., 2026), and their ability to uptake a number of desferrioxamine analogues has also been reported (D’Onofrio *et al*., 2010; Smahajcsik *et al*., 2026).

The findings have significant implications for sustainable pathogen control in aquaculture. The identification of a naturally occurring microbial consortium with potent, constitutively expressed anti-pathogenic properties offers a promising alternative to antibiotics. The dominant role of *Vreelandella alkaliphila* and *Alteromonas macleodii* in the inhibitory microbiome suggests their potential as core members of a rationally designed probiotic consortium. Understanding the ecological principles driving these interactions, such as iron competition, can inform strategies for microbiome engineering to enhance disease resistance in aquaculture systems. Future research should focus on validating these inhibitory mechanisms *in vivo* and assessing the stability and efficacy of such consortia under aquaculture conditions.

In conclusion, this multi-omics investigation provides a comprehensive view of the molecular mechanisms underlying pathogen inhibition by an *Isochrysis galbana* associated microbiome. We demonstrate that the production of siderophores, primarily by *Vreelandella alkaliphila,* leading to iron competition, is a key strategy for suppressing *V. anguillarum* in this system. This work not only advances our understanding of chemical ecology in marine microbial communities but also provides a scientific basis for developing novel, sustainable disease control strategies in the form of multi-strain probiotics for the aquaculture sector.

## ACKNOWLEDGEMENTS

The authors thank Mikael Lenz Strube for valuable advice regarding the ONT metagenomics sequencing, and Jette Melchiorsen for assistance with RNA extraction. This study has received funding from the European Union’s Horizon 2020 research and innovation programme under grant agreement no. 101000392 (MARBLES) and the Netherlands Organisation for Scientific Research NWO (NWO-XL programme OCENW.XL21.XL21.088 to M.H.M.). The Thermo Orbitrap Tribrid IQ-X LC-MS system was funded by the UK BBSRC ALERT-22 Scheme award BB/X019802/1.

The output reflects only the author’s view, and the Research Executive Agency (REA) cannot be held responsible for any use that may be made of the information contained therein.

## DATA AVAILABILITY

Raw 16S amplicon sequencing, metagenomics, and metatranscriptomics sequencing data have been deposited in the European Nucleotide Archive (ENA) under BioProject accession PRJEB110475. Raw metabolomics data have been deposited in the MassIVE repository under accession MSV000101136.

## CONFLICT OF INTEREST

The authors declare no conflicts of interest.

## Notes

### Competing Interest Statement

The authors have declared no competing interest.

