## Supplementary Information for "Multi-omics reveals mechanisms behind pathogen inhibition by a microalgal microbiome"

^#^ contributed equally

Running title: Multi-omics of microbiome mediated pathogen control

Marnix H. Medema,, +31 654758321

ORCID numbers

Dóra Smahajcsik: 0000-0002-6103-7283

Robert A. Koetsier: 0000-0002-4477-5401

Emmanuel Tope Oluwabusola: 0000-0003-3153-4323

Matilde Emídio Almeida: 0009-0008-3960-9397

Line Roager: 0000-0002-7033-7309

Scott A Jarmusch: 0000-0002-1021-1608

Morten Dencker Schostag: 0000-0001-9221-1398

Joseph Nesme: 0000-0003-1929-5040

Marcel Jaspars: 0000-0002-2426-6028

Lone Gram: 0000-0002-1076-5723

Marnix Medema: 0000-0002-2191-2821

**Table S1.** Primers with 30 unique octametric barcodes used for amplification of the V3-V4 region of the 16S rRNA gene in DNA extracted from samples of the native and inhibitory *Isochrysis galbana* microbiome (Klindworth *et al.*, 2013).

| **Barcode no.** | **Barcode** | **Forward primer (5’-3’) (w/ barcode)** | **Reverse primer (5’-3’) (w/ barcode)** |
| --- | --- | --- | --- |
| 1 | TTTTAATC | TTTTAATCCCTACGGGNGGCWGCAG | TTTTAATCGACTACHVGGGTATCTAATCC |
| 2 | ATAATTAG | ATAATTAGCCTACGGGNGGCWGCAG | ATAATTAGGACTACHVGGGTATCTAATCC |
| 3 | ACCAAATT | ACCAAATTCCTACGGGNGGCWGCAG | ACCAAATTGACTACHVGGGTATCTAATCC |
| 4 | CTTATCAA | CTTATCAACCTACGGGNGGCWGCAG | CTTATCAAGACTACHVGGGTATCTAATCC |
| 5 | TGATCATT | TGATCATTCCTACGGGNGGCWGCAG | TGATCATTGACTACHVGGGTATCTAATCC |
| 6 | AGAATCTA | AGAATCTACCTACGGGNGGCWGCAG | AGAATCTAGACTACHVGGGTATCTAATCC |
| 7 | TCAAGAAA | TCAAGAAACCTACGGGNGGCWGCAG | TCAAGAAAGACTACHVGGGTATCTAATCC |
| 8 | ATCGAAAT | ATCGAAATCCTACGGGNGGCWGCAG | ATCGAAATGACTACHVGGGTATCTAATCC |
| 9 | ACATTTAC | ACATTTACCCTACGGGNGGCWGCAG | ACATTTACGACTACHVGGGTATCTAATCC |
| 10 | TAGAAAAC | TAGAAAACCCTACGGGNGGCWGCAG | TAGAAAACGACTACHVGGGTATCTAATCC |
| 11 | TTATCACC | TTATCACCCCTACGGGNGGCWGCAG | TTATCACCGACTACHVGGGTATCTAATCC |
| 12 | AATAGGGT | AATAGGGTCCTACGGGNGGCWGCAG | AATAGGGTGACTACHVGGGTATCTAATCC |
| 13 | ATTGCTGA | ATTGCTGACCTACGGGNGGCWGCAG | ATTGCTGAGACTACHVGGGTATCTAATCC |
| 14 | TGAGTTCT | TGAGTTCTCCTACGGGNGGCWGCAG | TGAGTTCTGACTACHVGGGTATCTAATCC |
| 15 | GGCTATTT | GGCTATTTCCTACGGGNGGCWGCAG | GGCTATTTGACTACHVGGGTATCTAATCC |
| 16 | CAAGAGAT | CAAGAGATCCTACGGGNGGCWGCAG | CAAGAGATGACTACHVGGGTATCTAATCC |
| 17 | GGAATACA | GGAATACACCTACGGGNGGCWGCAG | GGAATACAGACTACHVGGGTATCTAATCC |
| 18 | AAGGCAAT | AAGGCAATCCTACGGGNGGCWGCAG | AAGGCAATGACTACHVGGGTATCTAATCC |
| 19 | ACAAAACG | ACAAAACGCCTACGGGNGGCWGCAG | ACAAAACGGACTACHVGGGTATCTAATCC |
| 21 | TTGAGTGA | TTGAGTGACCTACGGGNGGCWGCAG | TTGAGTGAGACTACHVGGGTATCTAATCC |
| 22 | GCTTCTGA | GCTTCTGACCTACGGGNGGCWGCAG | GCTTCTGAGACTACHVGGGTATCTAATCC |
| 23 | GGCAAGAT | GGCAAGATCCTACGGGNGGCWGCAG | GGCAAGATGACTACHVGGGTATCTAATCC |
| 24 | GTGCTTTC | GTGCTTTCCCTACGGGNGGCWGCAG | GTGCTTTCGACTACHVGGGTATCTAATCC |
| 30 | GCTTGGTT | GCTTGGTTCCTACGGGNGGCWGCAG | GCTTGGTTGACTACHVGGGTATCTAATCC |

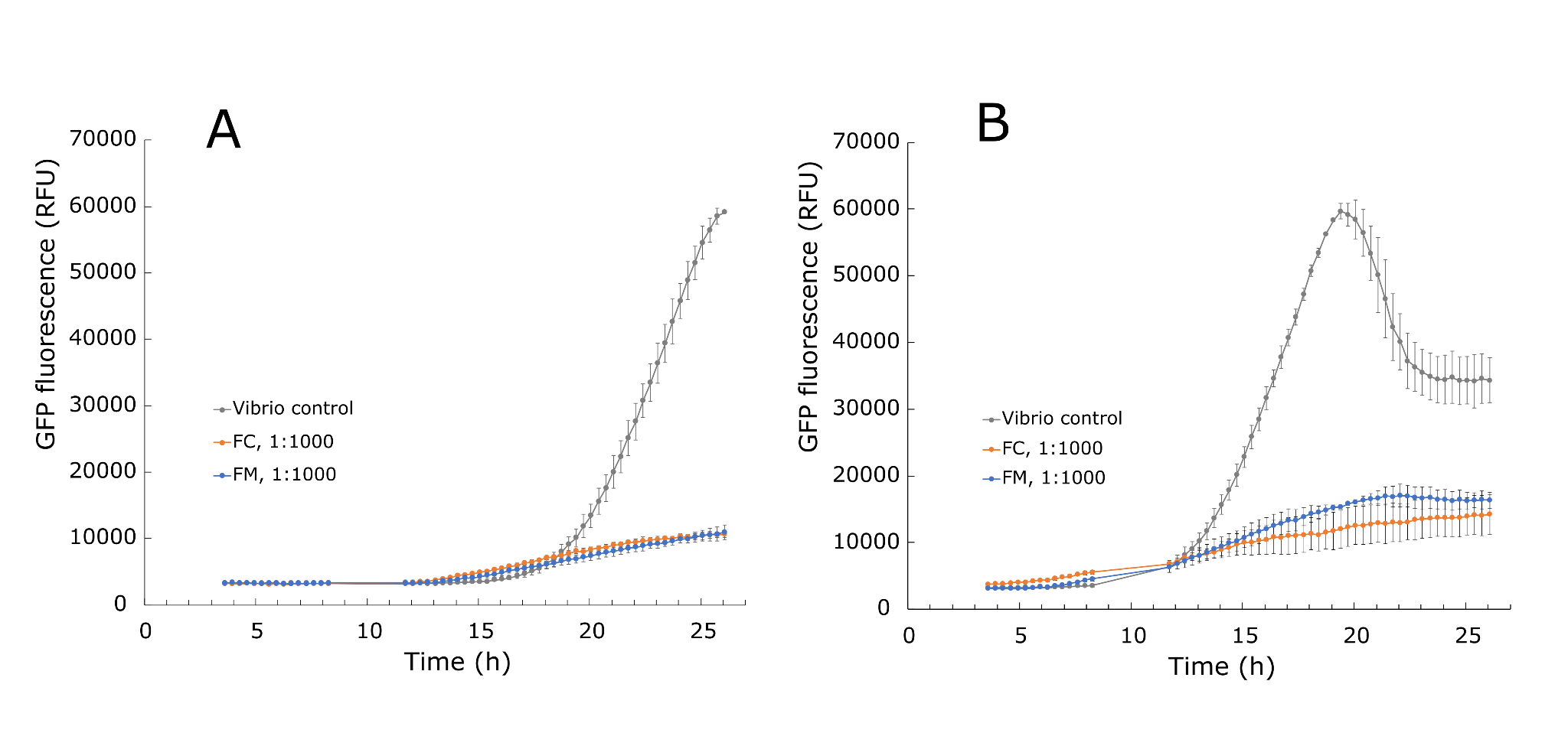

**Figure S1.** Inhibition of *Vibrio anguillarum* 90-11-286_gfp by FC and FM fractions of the *I. galbana* microbiome at a 1:1000 pathogen-to-microbiome ratio. GFP fluorescence of *V. anguillarum* co-cultured with the full culture (FC) and filtered microbiome (FM) fractions at **(A)** 2 log CFU/mL and **(B)** 4 log CFU/mL initial *Vibrio* inoculation density (n=4). A no-microbiome control is included for reference (n=3). Data represent mean ± SD.

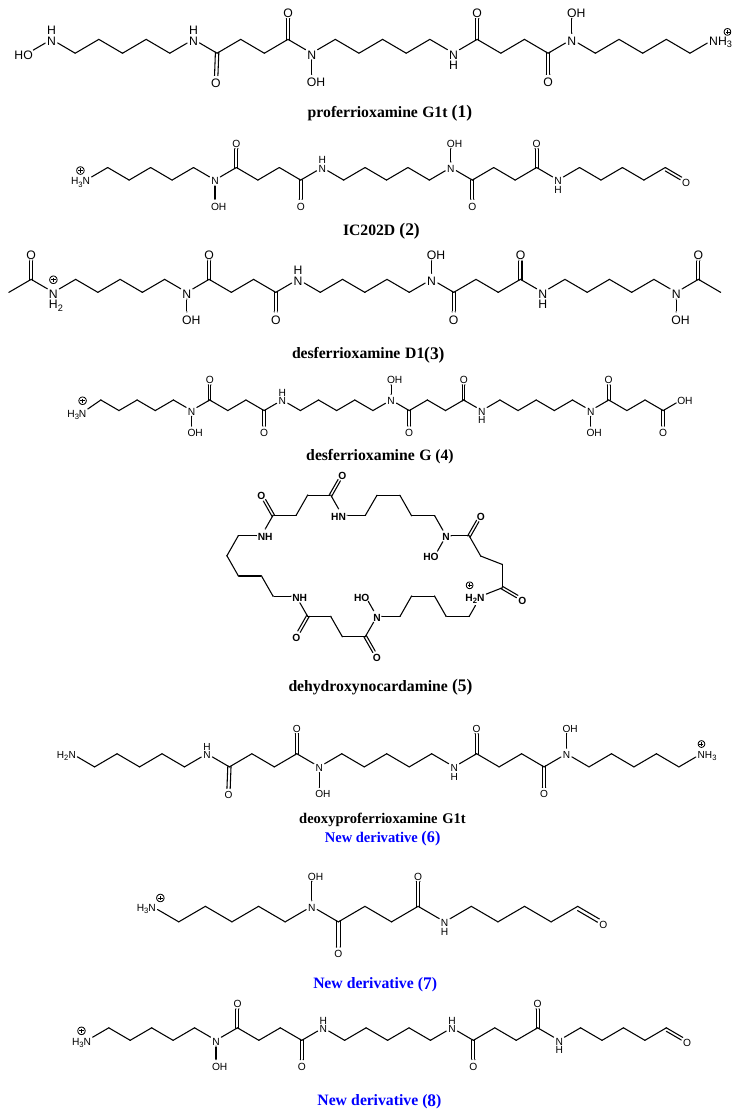

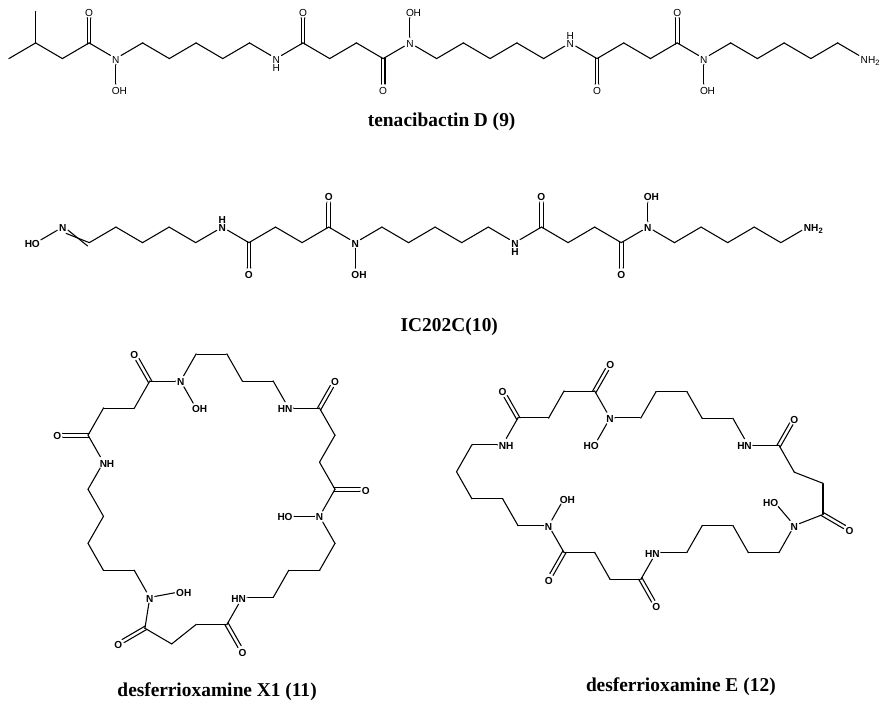

**Figure S2.** Putative compounds dereplicated from the VFM2 and VFM4 extracts.

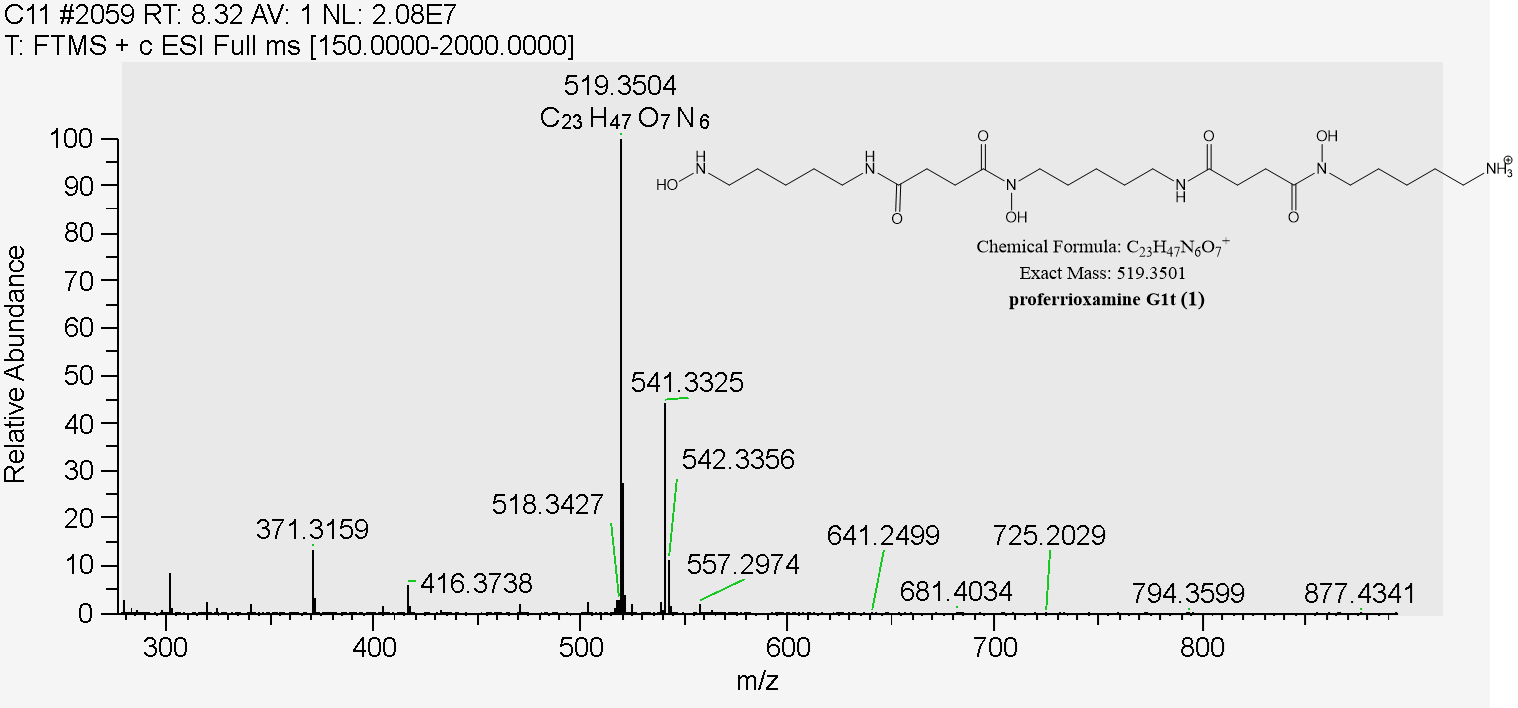

**Figure S3.** (+)-HR-ESIMS spectrum of **1**.

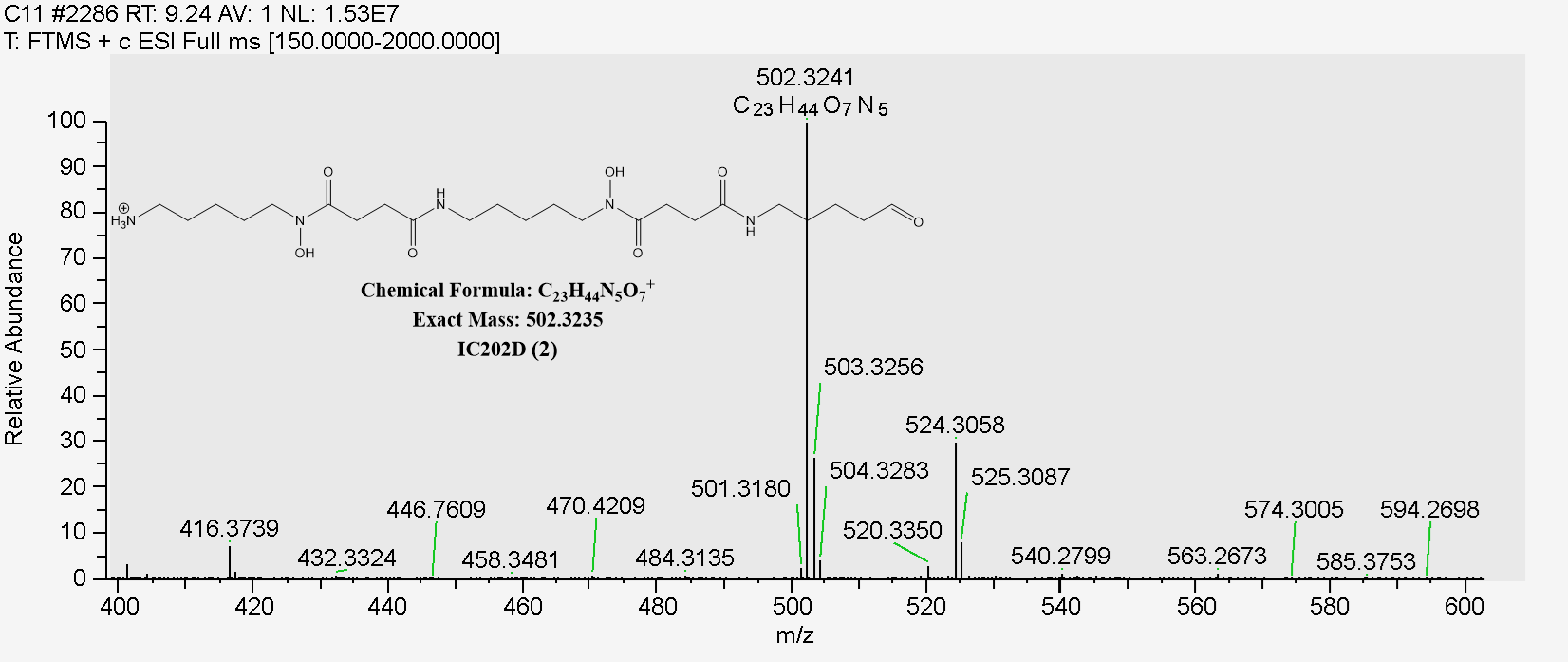

**Figure S4.** (+)-HR-ESIMS spectrum of **2**.

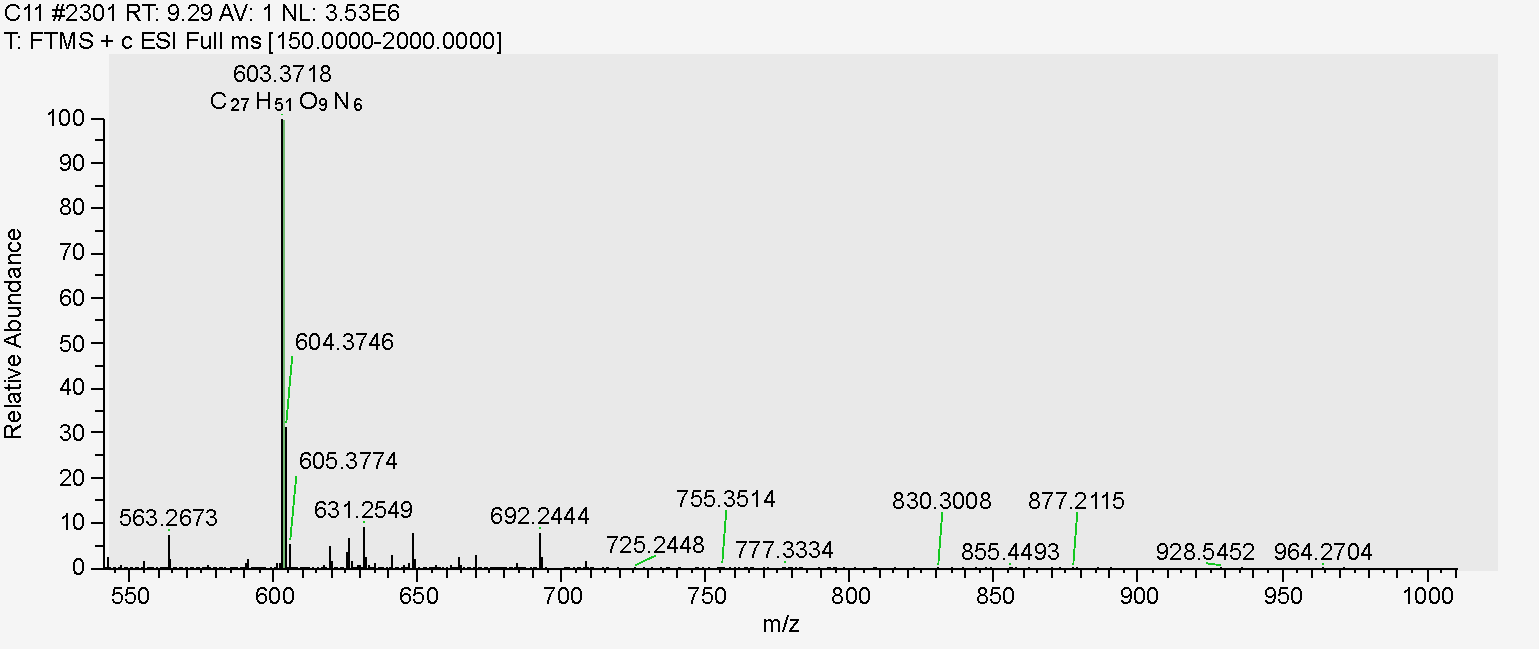

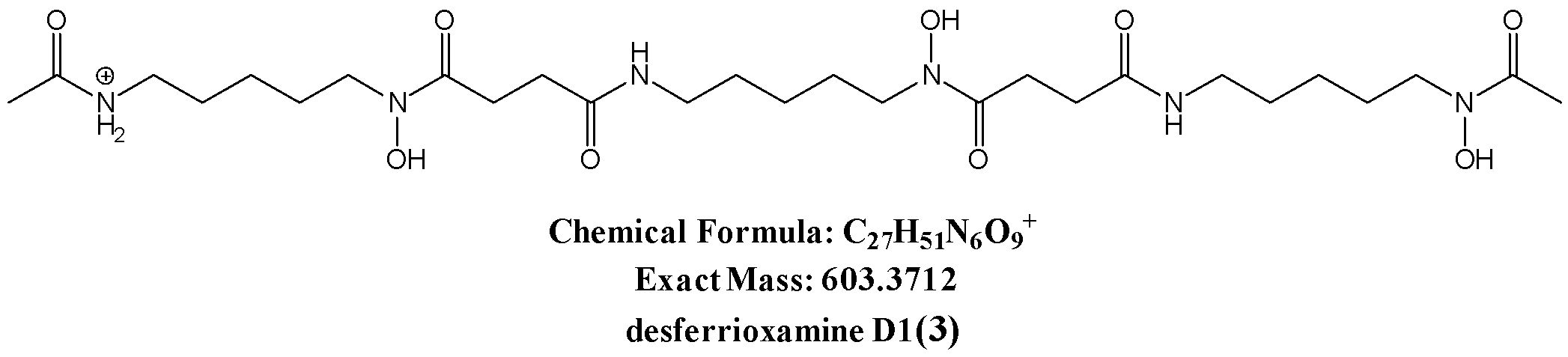

**Figure S5.** (+)-HR-ESIMS spectrum of **3**.

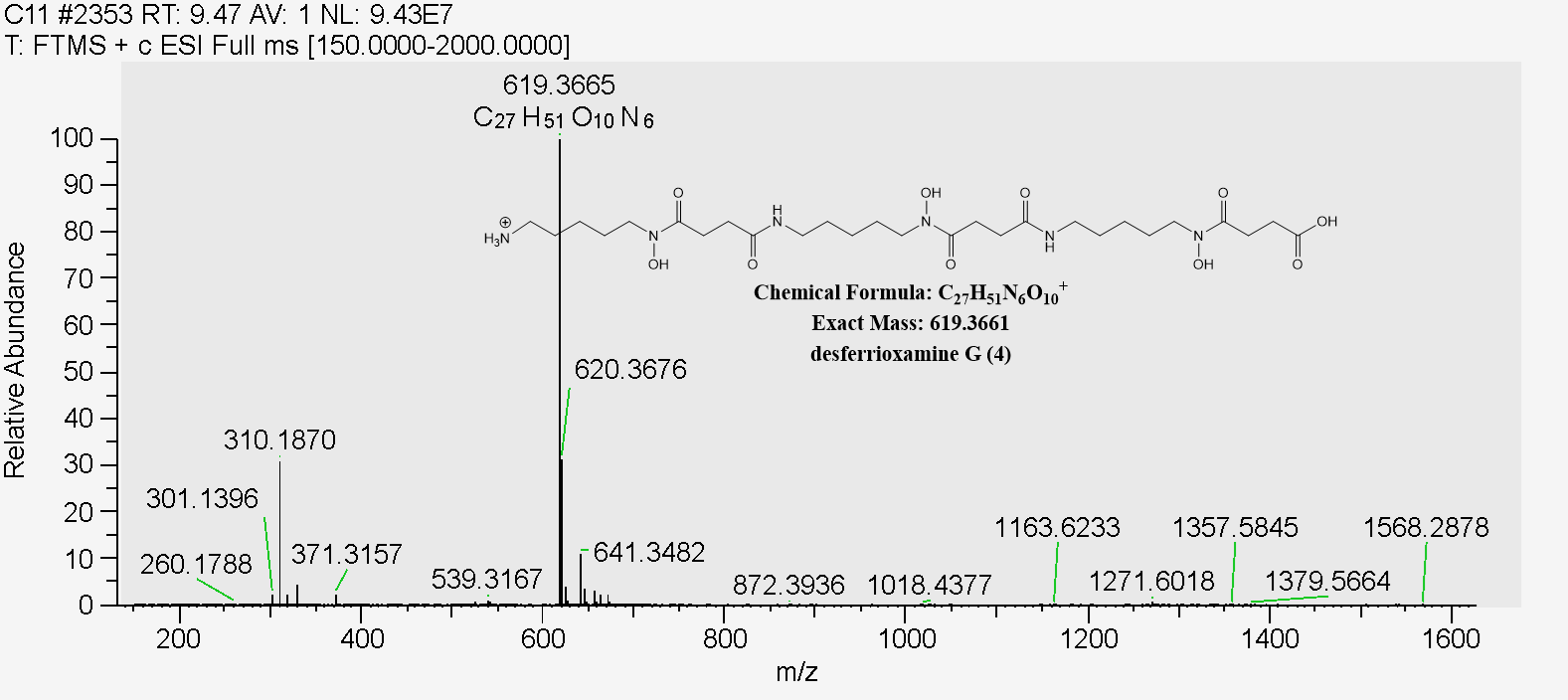

**Figure S6.** (+)-HR-ESIMS spectrum of **4.**

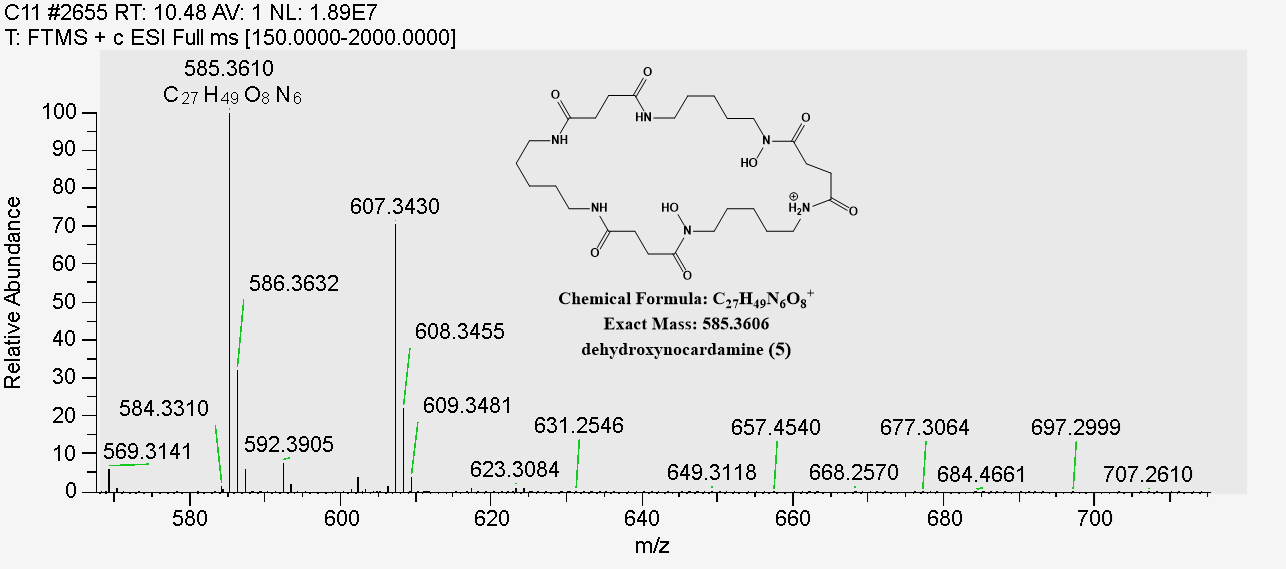

**Figure S7.** (+)-HR-ESIMS spectrum of **5.**

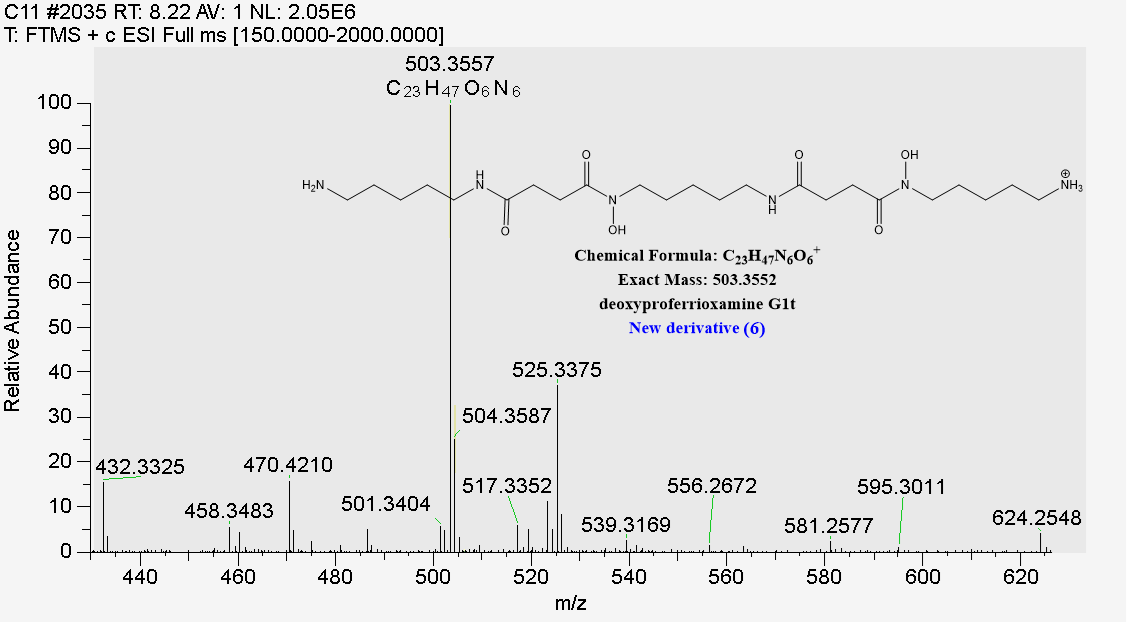

**Figure S8.** (+)-HR-ESIMS spectrum of **6.**

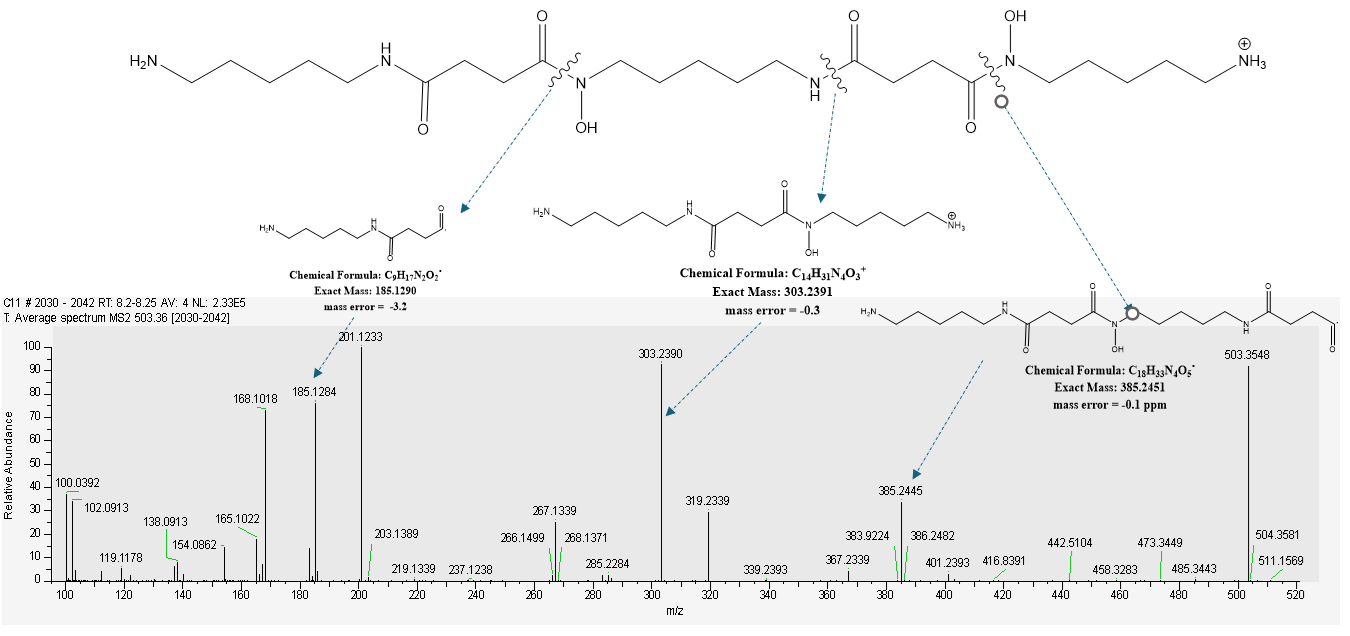

**Figure S9.** Fragment ions observed in HR-MS/MS spectrum for **6.**

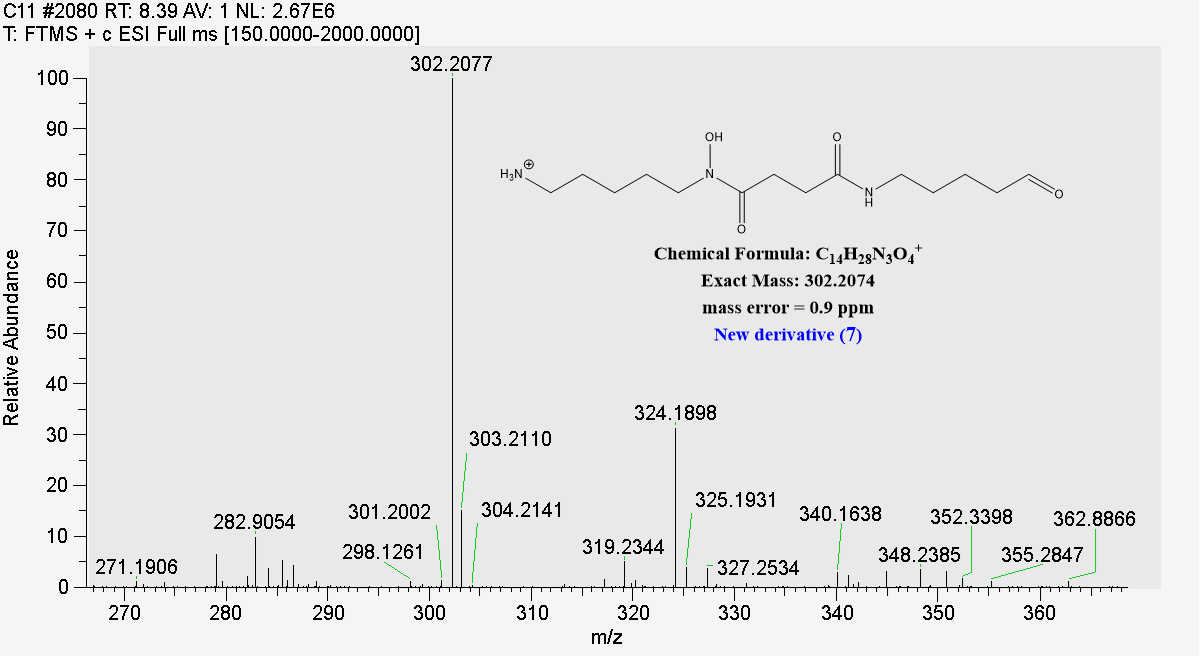

**Figure S10.** (+)-HR-ESIMS spectrum of **7.**

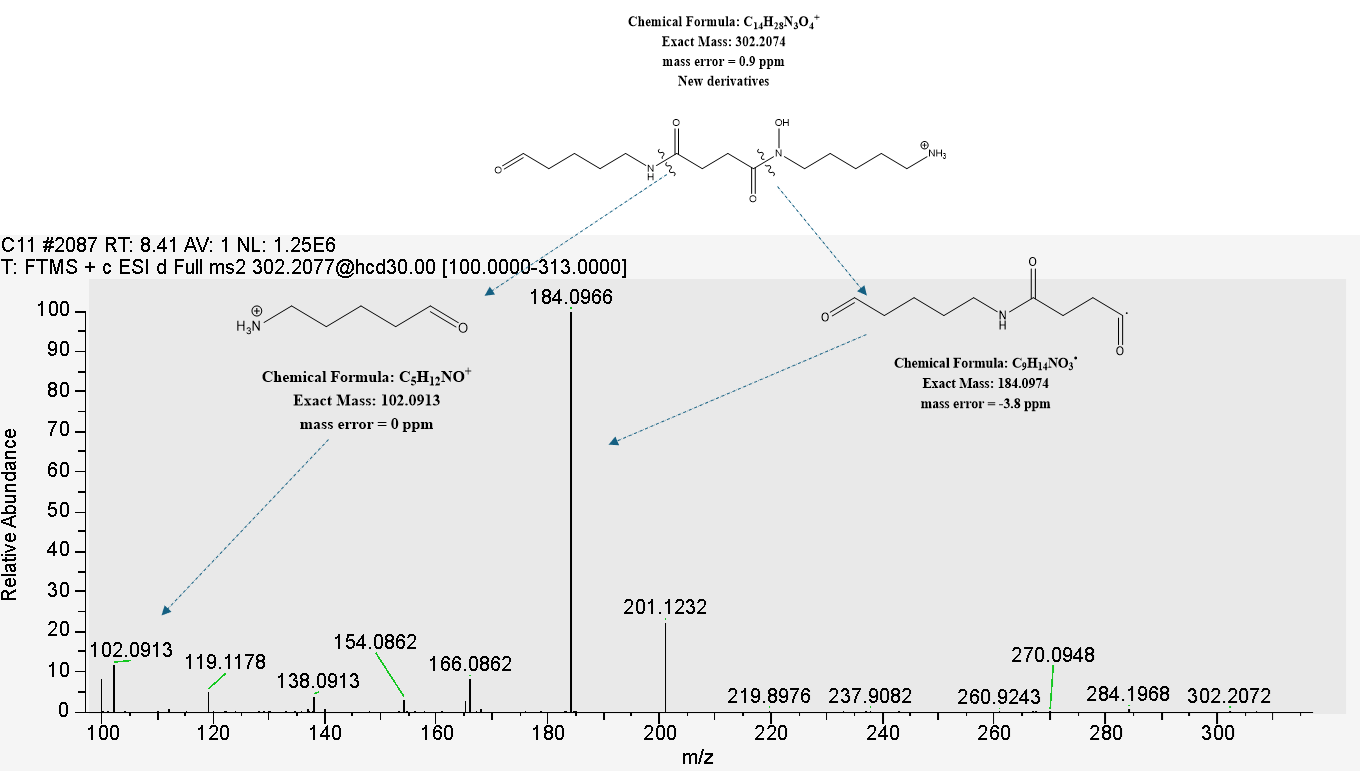

**Figure S11.** Fragment ions observed in HR-MS/MS spectrum for **7**.

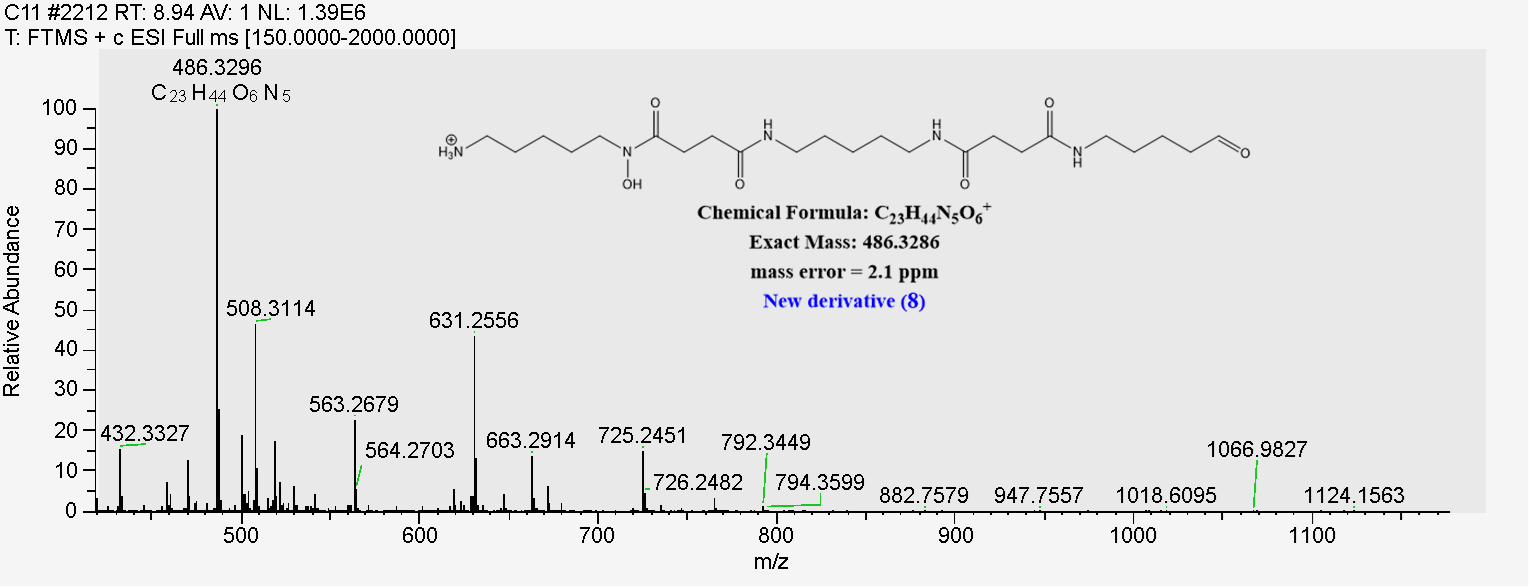

**Figure S12.** (+)-HR-ESIMS spectrum of **8.**

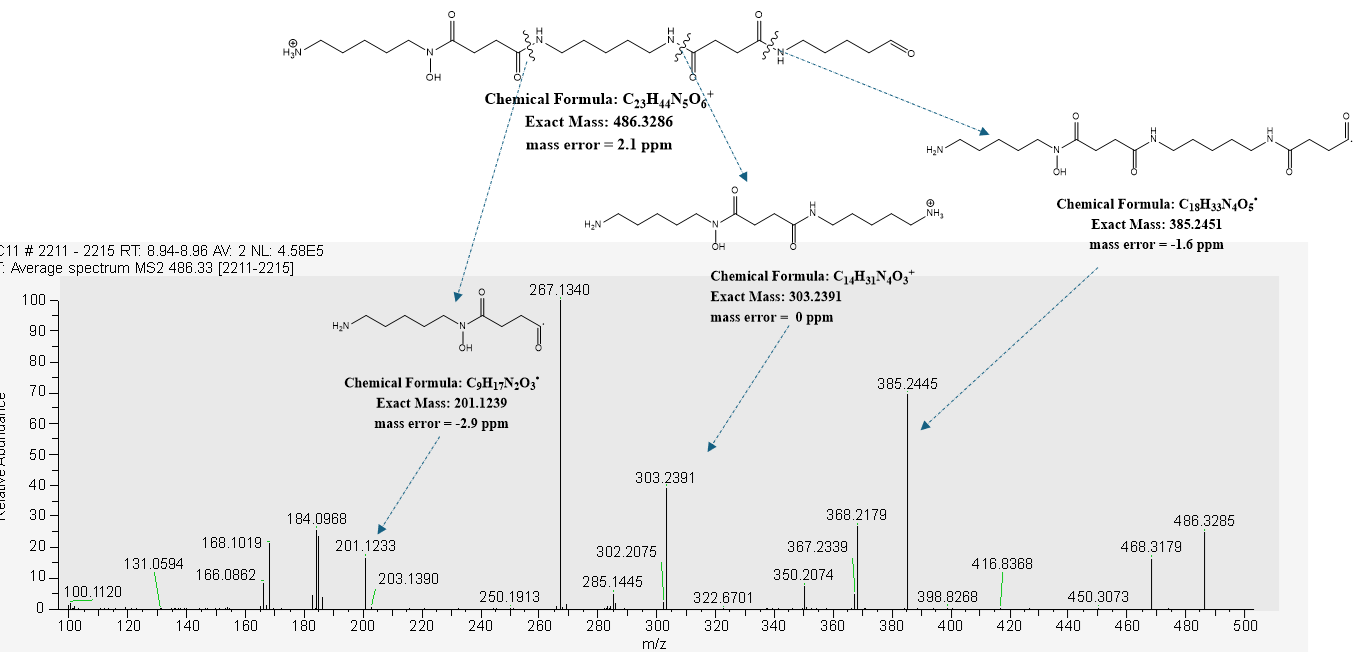

**Figure S13**. Fragment ions observed in HR-MS/MS spectrum for **8**.

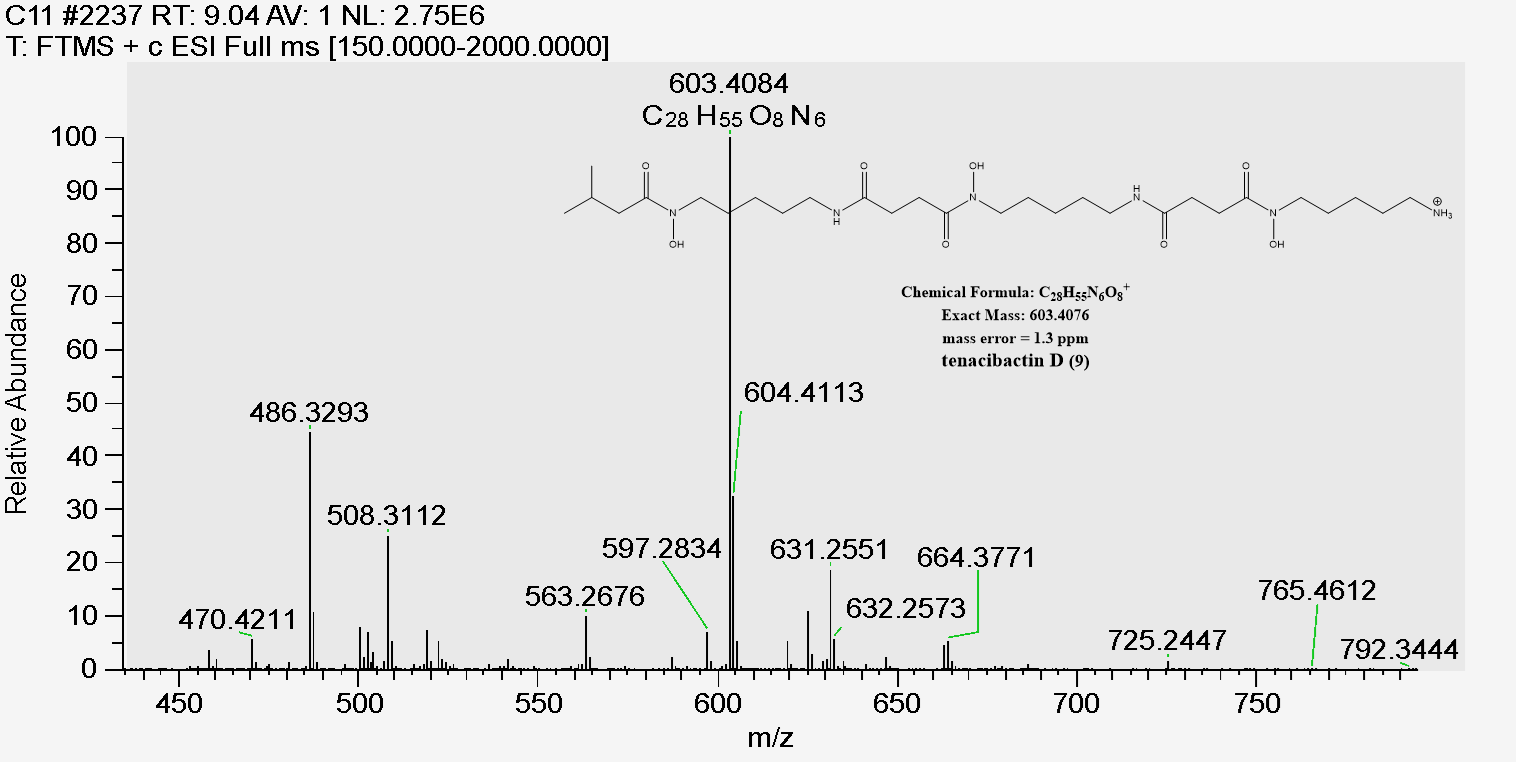

**Figure S14.** (+)-HR-ESIMS spectrum of **9.**

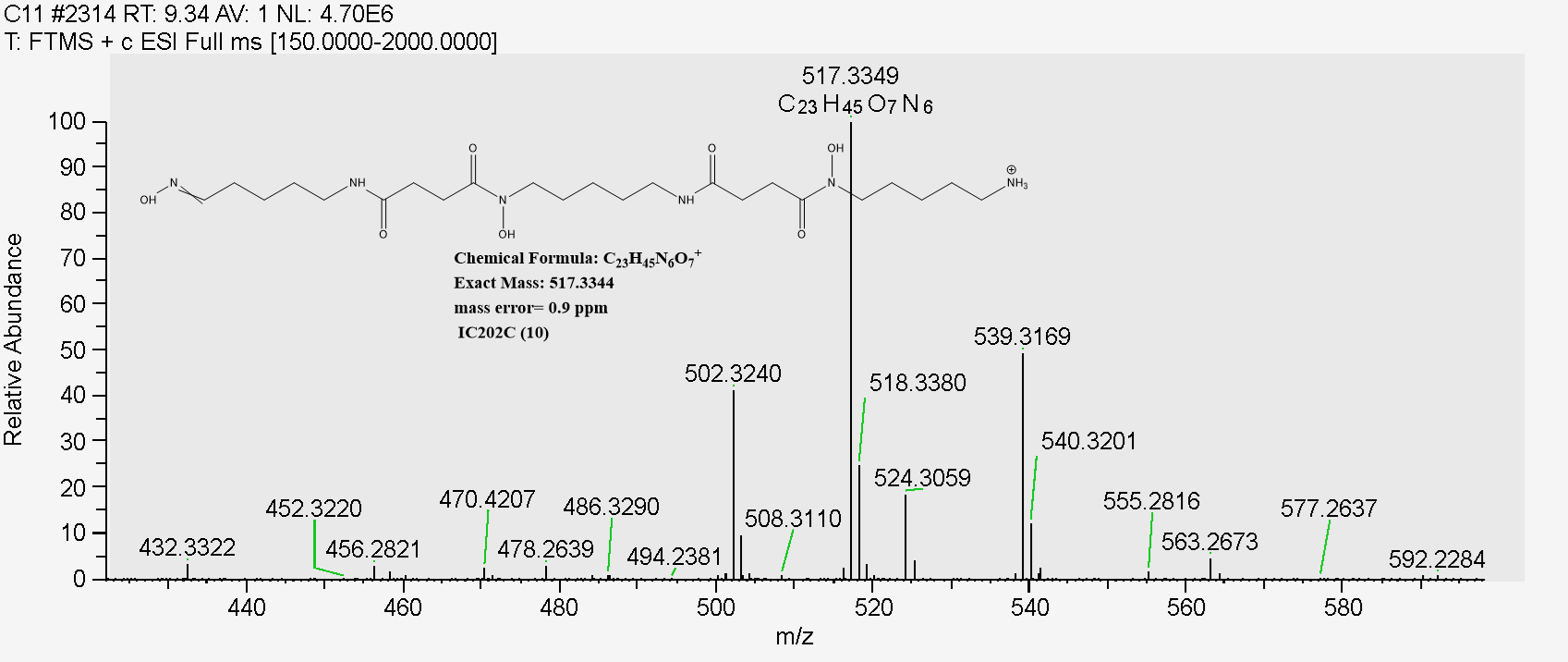

**Figure S15.** (+)-HR-ESIMS spectrum of **10.**

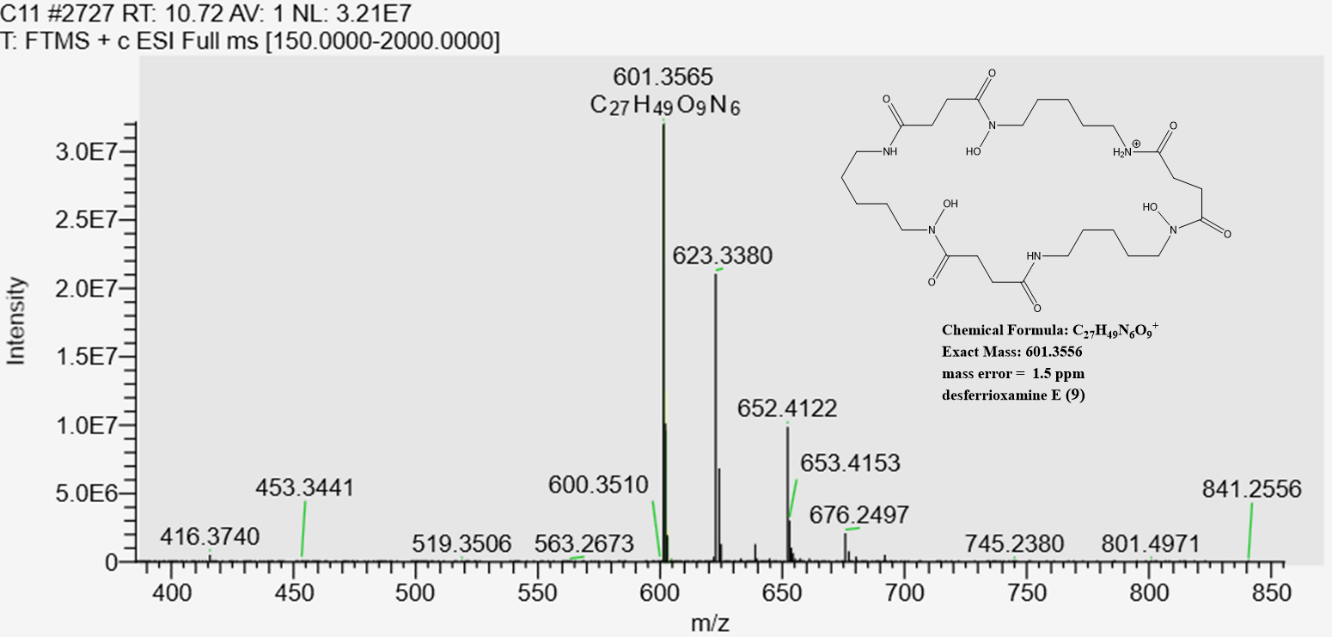

**Figure S16.** (+)-HR-ESIMS spectrum of **11.**

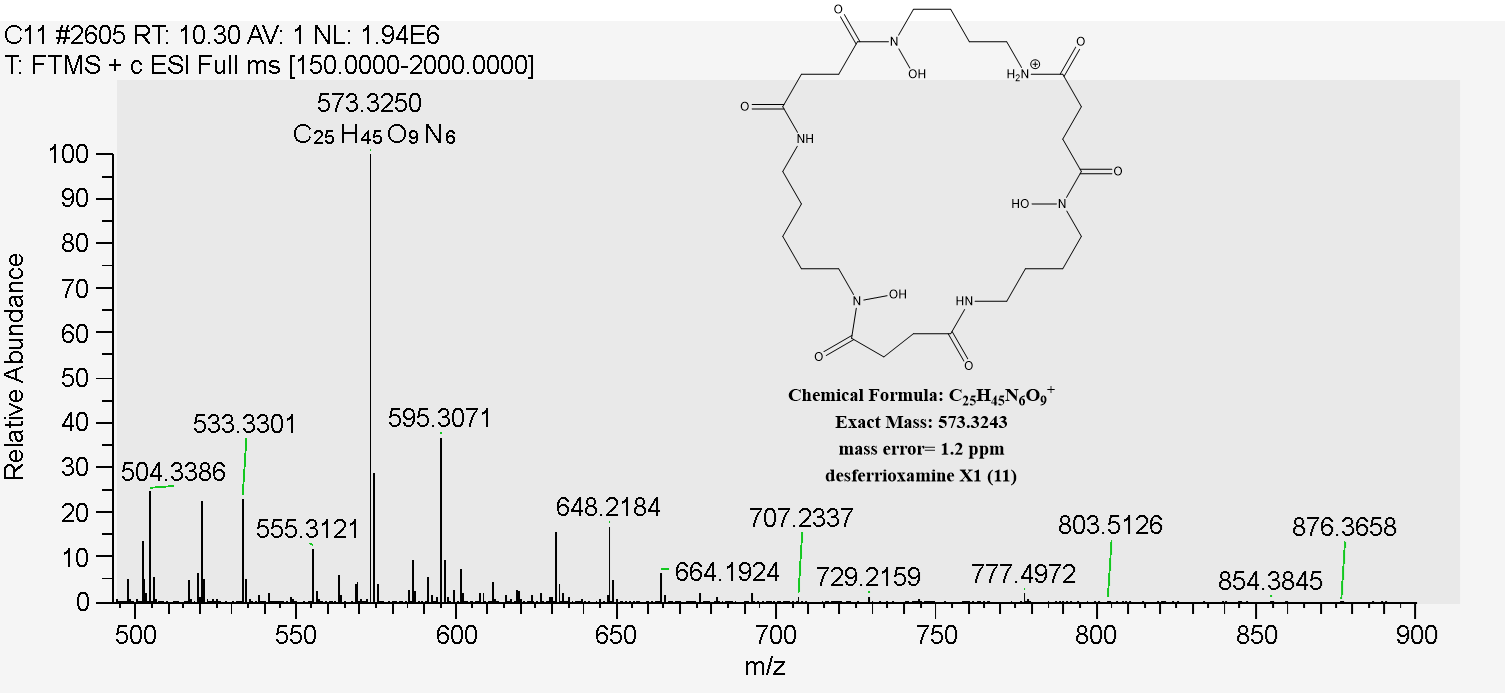

**Figure S17.** (+)-HR-ESIMS spectrum of **12.**

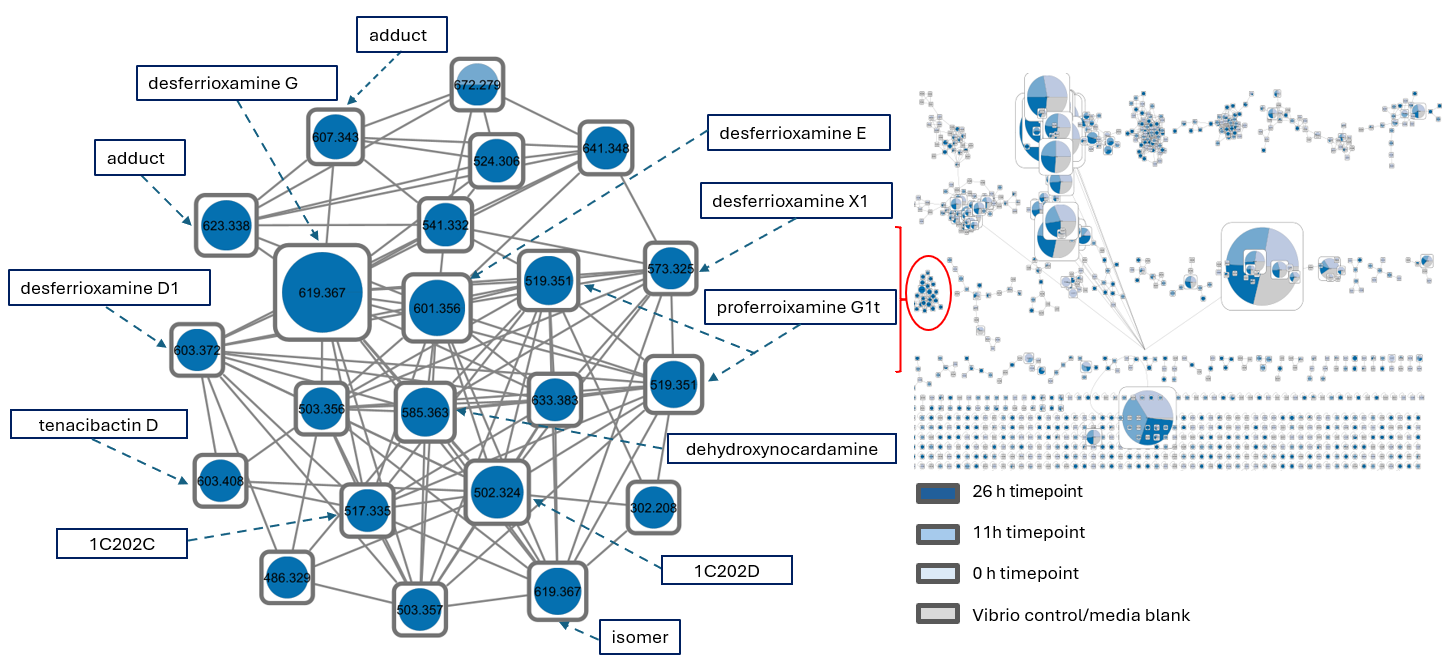

**Figure S18.** Feature-based MN analysis of Vibrio+FM2 (VFM2) extract exhibiting full inhibition at 26hrs.

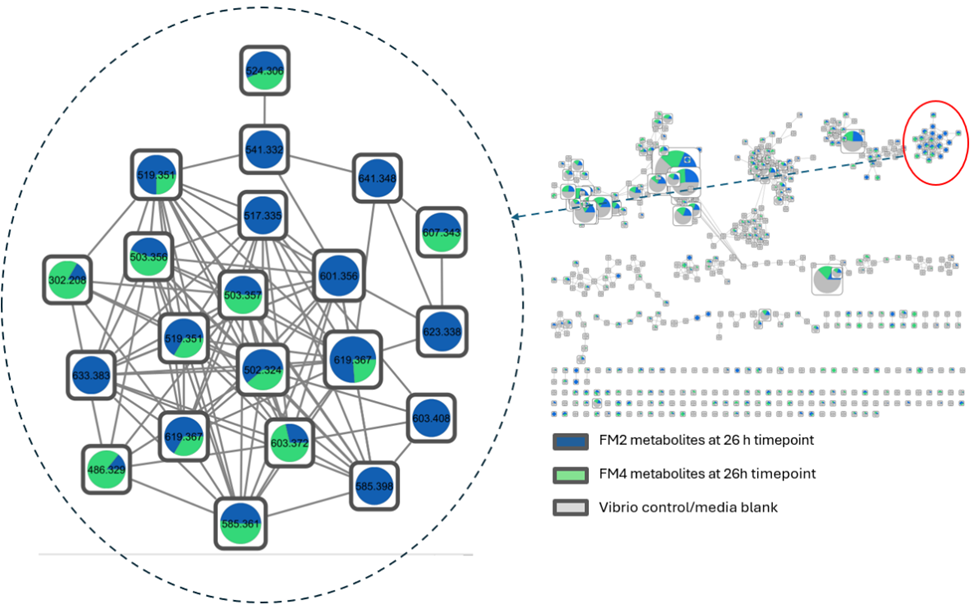

**Figure S19**. Feature-based Molecular Networking analysis comparing Vibrio + FM4 (VFM4; partial inhibition, marginal pathogen growth) and Vibrio + FM2 (VFM2; full inhibition) extracts exhibiting marginal pathogen growth (partial inhibition) at 26hrs.
